# A non-invasive approach to study localized forces in 3D epithelia

**DOI:** 10.64898/2026.04.01.715811

**Authors:** N. Gouirand, M. Ibrahimi, C. Valotteau, B. Lecouffe, A. Le Bivic, D. Massey-Harroche, F. Rico, M. Merkel, D Delacour, E Bazellières

## Abstract

Epithelial tissue integrity and remodelling rely on mechanical coupling between cytoskeletal organization and adhesion complexes, but their orchestration in three-dimensional (3D) epithelia remains unresolved. Here, we introduced a non-invasive strategy to locally probe forces within 3D epithelial tissues. We engineered elastic polyacrylamide (PAAm) microbeads with cell-like size and mechanical properties, enabling their integration into tissues. PAAm microbeads were spontaneously engulfed by the tissue, establishing an intrinsic interface. This allowed bead deformation to be directly correlated with local cytoskeletal architecture and adhesion organization, visualized through high-resolution imaging and quantitative 3D computational reconstruction. Using this approach, we demonstrated that localized mechanical perturbations trigger cytoskeletal remodelling while preserving tissue polarity. We further identified extracellular matrix composition as a key determinant of bead-tissue interactions. At the bead–cell interface, cells assembled tension-bearing focal adhesions and actin stress fibers. Quantitative analysis of bead deformation revealed sub-cellular scale patterns of cell-bead interaction forces, which correlated with the localization of cytoskeletal elements. Our findings extend previous traction force microscopy observations to 3D, revealing coordinated, spatially heterogeneous forces specifically at the cell–bead interface. Taken together, this enabled us to dissect the interplay between cellular processes and tissue mechanics in a non-invasive way in 3D model systems.

## Introduction

The development, organization, and homeostasis of epithelial tissues are governed by the multiscale integration of biochemical signalling and mechanical interactions across molecular, cellular, and tissue levels. In polarized epithelia, tissue architecture is established through the precise spatial organization of adhesion complexes and cytoskeletal networks along the apico–basal axis, generating a dynamic and mechanically responsive system that continuously interacts with compliant environments. At the core of this organization, basal focal adhesions (FAs), lateral adherens junctions (AJs), and apical tight junctions (TJs) collectively establish epithelial polarity while ensuring mechanical coupling both between neighbouring cells and with the extracellular matrix (ECM)[1]. These adhesive complexes are interconnected through the actin cytoskeleton, which provides structural integrity while enabling adaptive remodelling during development, regeneration, and disease progression. Mechanical forces transmitted through actomyosin networks and adhesion complexes are continuously converted into biochemical signals, allowing epithelial cells to dynamically regulate their shape, polarity, and collective behaviour in response to their physical environment [2–5]. This mechano-biochemical coupling is central to epithelial function, yet its integration across subcellular compartments and across multiple length scales remains poorly understood.

In columnar epithelia, distinct actin architectures sustain specialized mechanical functions. At the apical domain, branched apical actin networks — nucleated by Arp2/3 and controlled by Rac and Cdc42 signalling — stabilize AJs and TJs by modulating membrane tension and trafficking dynamics [6–10]. These networks generate protrusive forces and are increasingly recognized as key regulators of mechanosensitive pathways, notably through Piezo1, which tunes actin remodelling in response to extracellular stiffness [11]. In parallel, contractile actin bundle systems, including the junctional actin belt and medial–apical fibers, generate tensile forces that drive apical constriction and epithelial folding. These morphogenetic processes require the coordinated production and transmission of forces across individual cells and tissues [5,12]. Actin bundles are structurally organized by crosslinkers such as fascin, fimbrin, and α-actinin, and their contractility is regulated by the RhoA–ROCK–myosin-II pathway [13]. At the tissue scale, AJs and TJs function not only as structural elements but also as active components for force transmission and mechanochemical signalling [14–17]. Moreover, emerging evidence further suggests isoform-specific contributions of non-muscle myosin-II to force generation and mechanical specialization with epithelial tissues [18]. During collective behaviours such as migration, branched actin networks promote protrusive activity through lamellipodia formation, whereas bundled actin structures reinforce FAs and intercellular junctions, enabling coordinated force propagation across multicellular assemblies [19–23].

Despite these advances, the mechanisms governing the regulation of force-generating cytoskeletal architectures within polarized 3D epithelial tissues interacting with compliant materials remain poorly understood. Current methodologies, particularly 3D Traction Force Microscopy (TFM), have provided important insights into cell-matrix force transmission. However, their broad application is fundamentally limited by high computational complexity, dependence on nonlinear finite element modelling, and the requirement for precisely defined material parameters that are often difficult to obtain in biologically heterogeneous matrices [24–26]. Moreover, these approaches rely on assumptions — including well-defined stress-free reference states and idealized boundary conditions — that are rarely met in living systems, thereby restricting their accuracy and robustness [27–29]. Critically, the mechanical coupling between cells and their surrounding matrix is frequently incomplete, especially in stiff environments where cellular forces fail to produce measurable matrix deformations [26]. Finally, the lack of independent *in vivo* benchmarks for force measurements remains a major obstacle to validation and quantitative interpretation [27]. Together, these limitations highlight a central unresolved challenge: the need of robust, quantitative, and physiologically relevant approaches to measure and interpret force generation within intact 3D epithelial systems.

To address these challenges, biohybrid force-sensing materials directly integrated into living tissues have recently emerged as powerful tools for probing multiscale mechanics *in situ*. By combining well-defined material properties with biological compatibility, these systems enable localized and quantitative measurements of cellular forces within confined and dynamically evolving environments [30]. Pioneering approaches based on oil microdroplets provided the first direct evidence of anisotropic, myosin-II-dependent stresses in 3D cell aggregates and tissues *in vivo* [31–33]. More recently, double-emulsion droplets have extended this strategy toward the measurement of osmotic pressures [33]. However, the mechanical response of oil droplets is strongly influenced by interfacial tension, limiting their use as quantitative reporters of hydrostatic stresses and volumetric deformations. To overcome this concern, hydrogel-based sensors, including alginate and <u>p</u>oly<u>a</u>cryl<u>am</u>ide (PAAm) microgels, have been developed as elastic biohybrid materials that deform in response to cellular forces. These systems enable the quantification of both isotropic and anisotropic stress components in 3D biological models, including spheroids, tumoroids, and developing tissues [34–36]. In addition, the incorporation of fluorescent tracers allows high-resolution 3D reconstruction of stress fields, although this typically requires advanced imaging and computational modelling pipelines [37,38].

In this study, we introduced a biohybrid material platform that allows non-invasive and spatially resolved force mapping within polarized epithelial 3D tissues. We engineered elastic PAAm microbeads as cell-mimetic force sensors with precise tunable stiffness and defined ECM coatings, thereby recapitulating key biochemical and mechanical features of the native microenvironment. Using Caco-2 cysts and intestinal organoids as model systems, we showed that—unlike conventional injection-based approaches—PAAm beads were actively engulfed by epithelial tissues in a coating-dependent manner. This led to the formation of mechanically coupled cell–material interfaces. In particular, collagen-I functionalization promoted robust adhesion and efficient tissue integration, enabling sustained physiologically relevant force transmission. At bead–cell interfaces, epithelial cells assembled tension-bearing FAs and organized actin stress fibers, revealing localized cytoskeletal remodelling and the emergence of active cortical stress without perturbing the global tissue polarity. By combining high-resolution confocal imaging with biochemical perturbations, and quantitative 3D reconstruction and deformation analysis, we established a direct link between bead deformation patterns and the underlying cytoskeletal architectures. These analyses uncovered spatially patterned bead-cell interaction forces, demonstrating that epithelial tissues deploy highly organized, anisotropic force distributions rather than uniform stresses. Together, our biohybrid force-sensing strategy provides a versatile materials-based framework for quantifying and mapping cellular force patterning in complex 3D tissues.

## Results

### Production and characterization of PAAm microbeads as force sensors

To enable quantitative, cell-resolved measurements of mechanical forces in 3D epithelial tissues, we developed a robust, scalable and highly tunable method for fabricating elastic polyacrylamide (PAAm) microbeads. Using a water-in-oil emulsion technique (Figure 1A), an aqueous precursor solution— containing acrylamide, bis-acrylamide crosslinker, and fluorescent reporters—was dispersed in a fluorinated oil phase with surfactant to prevent droplet coalescence. Emulsification was performed via repeated syringe extrusion through a 25G needle, producing spherical hydrogel microbeads (Figure 1A). To verify the incompressibility of the PAAm beads, we measured their lateral deformation while the AFM tip applied a force to the top (Supplementary Figure 1A). These measurements indicated a Poisson ratio of 0.34 ±0.17 (Supplementary Figure 1B), consistent with previously reported values [39,40]. These AFM measurements confirmed that the PAAm beads are nearly incompressible. Furthermore, mechanical tunability was achieved by adjusting the bis-acrylamide-to-acrylamide ratio, allowing precise control of bead stiffness across a physiologically relevant range (0.6 to 12 kPa).

**Figure 1:**
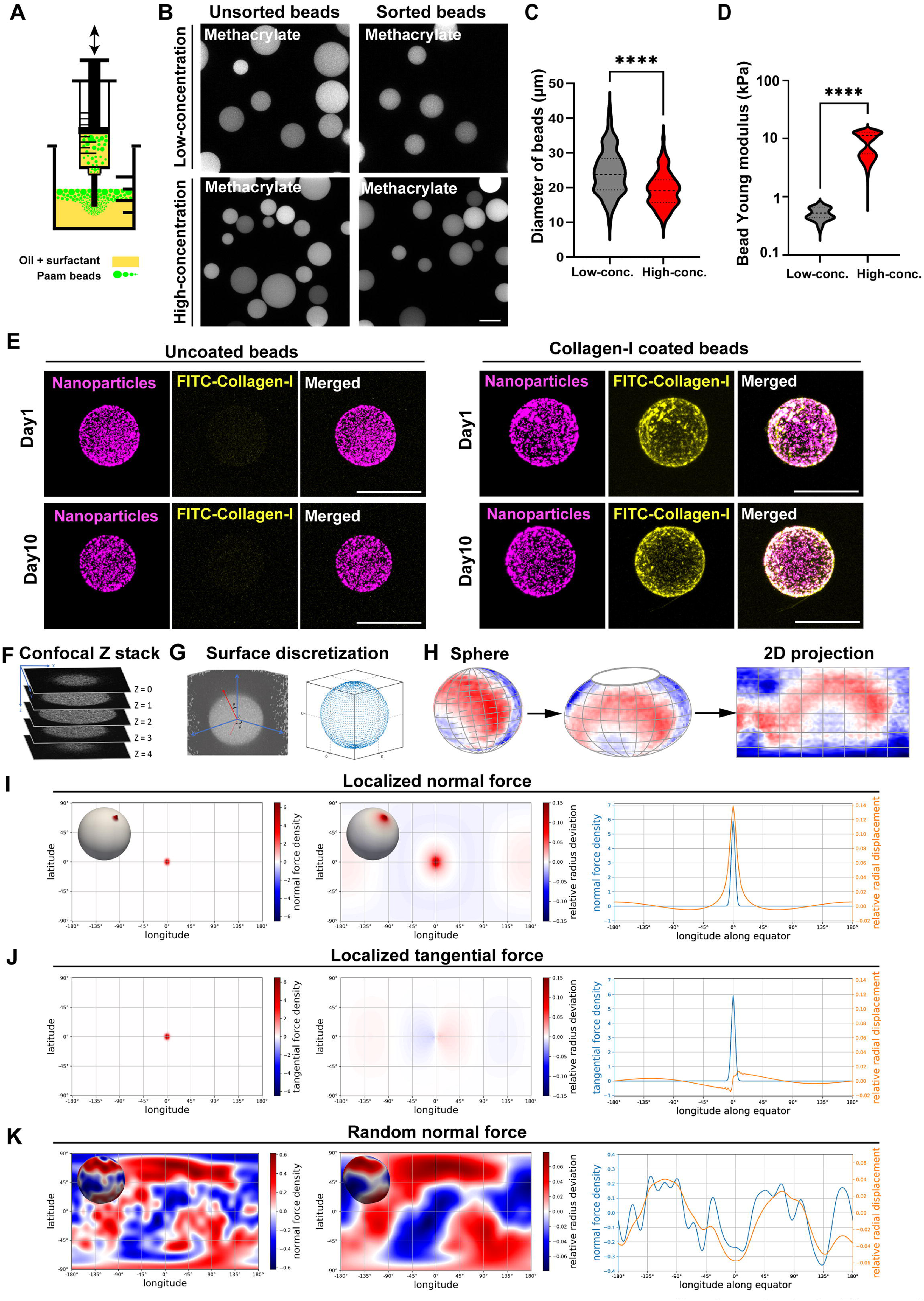
Production and characterization of deformable polyacrylamide (PAAm) beads. **A,** Schematic illustration of the procedure used for producing PAAm beads (green). A water-in-oil emulsion (yellow) was generated by passing the mixture through a syringe. **B**, Confocal analysis of the diameter of fluorescein-O-methacrylate PAAm beads as a function of bis-acrylamide crosslink concentration (low-concentration beads = 22 µl/ml, or high-concentration beads = 80 µl/ml of a 2% bis-acrylamide solution). Scale bar, 20μm. **C**, Statistical analysis of bead diameter based on bis-acrylamide concentration. Mean diameter of low-concentration beads = 24.41±6.52µm (mean±SD), of high-concentration beads = 19.4±4.99µm. n (low-concentration) = 124 beads, n (high-concentration) = 96 beads. Mann-Whitney test, ****p<0.001. **D**, Statistical analysis of the Young’s modulus of the PAAm beads. Mean Young’s modulus of low-concentration beads = 788±337Pa (mean±SD), of high-concentration beads = 8483±3745Pa. n (low-concentration) = 21 beads, n (high-concentration) = 21 beads. Mann-Whitney test, ****p<0.001. **E**, Confocal analysis of the distribution of far-red carboxylate-modified nanoparticles in PAAm beads, with or without FITC-collagen-I coating, after 1 or 10 days in PBS solution. Scale bar, 20μm. **F-H**, Workflow for 3D PAAm bead reconstruction. Sequential steps include z-stack image acquisition (**F**), coordinate transformation and alignment (**G**), mapping of the PAAM bead surface onto a spherical reference frame and projection into a 2D representation (**H**). (**I-K**) FEM simulations to probe the deformation patterns resulting from specific force patterns applied to the bead surface (see Methods for details). (**I**) A localized force applied normally to the bead surface (left) resulted in a localized radial displacement over a slightly broader region (middle). A plot of both along the equator allows detailed comparison (right). (**J**) A localized force applied tangentially to the bead surface (left) resulted in minimal radial displacement of the bead surface (middle), as confirmed by the plot along the equator (right). (**K**) A random pattern of normal forces applied to the bead surface (left) produced a radial displacement pattern that was essentially a blurred version of the applied force pattern (middle), confirmed by a plot along the equator (right).

To support high-resolution 3D reconstruction and quality control, we implemented dual fluorescent labelling strategies: homogeneous labelling with fluorescein-O-methacrylate (Figure 1B), or combined labelling with fluorescein-O-methacrylate and far-red carboxylate-modified nanoparticles (Supplementary Figure 1C). These approaches provided complementary insights into bead geometry and structural integrity.

We initially observed polydispersity in bead size (Supplementary Figure 1C), which we addressed by using sequential size filtration with mesh strainers to isolate cell-scale beads (Figure 1B). Post-filtration image analysis revealed that beads fabricated with a low crosslinker concentration (22 µl/ml of a 2% bis-acrylamide solution) had a mean diameter of 24.41 ± 6.52 µm, while those with a high crosslinker concentration (80 µl/ml) were slightly smaller, with a mean diameter of 19.40 ± 4.99 µm (Figure 1C). To quantify bead stiffness, we performed Atomic Force Microscopy (AFM) analysis, which revealed that the beads are purely elastic (Supplementary Figure 1B) and that low-crosslinker beads were softer, with a Young’s modulus of 788 ± 337 Pa (mean± SD), whereas highly crosslinked beads were stiffer, with a Young’s modulus of 8.48 ± 3.74 kPa (Figure 1D). Notably, this stiffness range spanned that of the native intestinal epithelium to colorectal tumour tissue [41], highlighting the physiological relevance of the system. For subsequent experiments, we selected stiff beads to model the carcinoma-associated microenvironment in Caco-2 cultures and in intestinal organoids, while soft beads were used to mimic the mechanical context of healthy intestinal organoids.

Furthermore, to create well-defined bio-interfaces, bead surfaces were functionalized with ECM proteins, such as collagen-I, using sulfo-SANPAH-mediated protein conjugation. Coating efficiency and spatial homogeneity were validated using FITC-labelled collagen-I, which produced a continuous and uniform fluorescent signal on the bead surface (Figure 1E). The collagen-I coating and cell culture conditions had no significant effect on bead shape (Supplementary Figure 1D-E). Comparison with uncoated beads confirmed no detectable disruption in the internal hydrogel structure (Figure 1E). Importantly, the collagen-I coatings remained stable for at least 10 days in cell culture medium (Supplementary Figure 1D-E), ensuring their suitability for long-term mechanical measurements in complex biological environments. Collectively, this fabrication pipeline yielded mechanically tunable, size-controlled PAAm microbeads with robust and customizable biofunctional surfaces, which could be used for mechanobiological interrogation in 3D epithelial systems.

Next, we attempted to directly measure *in situ* forces by inferring cell-bead interaction forces from the PAAm bead deformation (see Methods for details). Yet, this approach was not feasible, as it would require imaging both the deformed and undeformed states of the same bead, which is currently experimentally unachievable. As a consequence, in this manuscript, we cannot make any statement about any force that is applied homogeneously and perpendicularly to the bead surface, since that would lead at most to a change in the bead size, which we cannot measure. However, it is still possible to get an idea about *variations* in the applied forces, since they would distort the bead shape away from spherical. To this end, we used simulations to probe the effects of specific force patterns applied to the bead surface. Given that PAAm behaves as a linearly elastic solid over a wide range of deformations [42], we reasoned that studying bead deformation in response to localized normal or tangential forces exerted on the bead surface would suffice. Complex force patterns could then be decomposed into localized forces, with the resulting deformation pattern being the superposition of individual responses to these localized forces. Using Finite-Element-Method (FEM) simulations (see Methods for details), we confirmed that a locally applied normal (i.e. pulling) force (Figure 1I, left panel) produced a localized tensile deformation (Figure 1I, middle panel). Specifically, the local radial deviation peaked near the force application point, albeit with a slightly broader distribution, while deformation elsewhere remained minimal (Figure 1I, right panel). In contrast, a localized tangential force induced a smaller deformation (Figure 1J). Therefore, we interpreted measured bead deformations primarily in terms of applied normal forces. When applying a complex pattern of normal forces applied randomly across the surface, we found that the resulting radial deviation closely resembled a slightly blurred version of the applied normal forces (Figure 1K). This aligned with the observation that the deformation response to a localized normal force is also broader (Figure 1I). Taken together, while we could not yet quantitatively compute precise forces on the bead surface, these results suggested that radial deformation serves as a reliable proxy for the spatial variations in the local normal forces applied to the bead surface.

### ECM surface chemistry determines bead integration into polarized epithelia

Next, we investigated how ECM functionalization modulates bead-tissue interactions within polarized 3D epithelial architectures. To maintain tissue integrity and avoid artefacts resulting from invasive manipulations, we developed a non-disruptive assay. In this assay, ECM-functionalized hydrogel beads were introduced directly into the culture medium, enabling spontaneous interaction with pre-established epithelial structures (Figure 2A).

**Figure 2:**
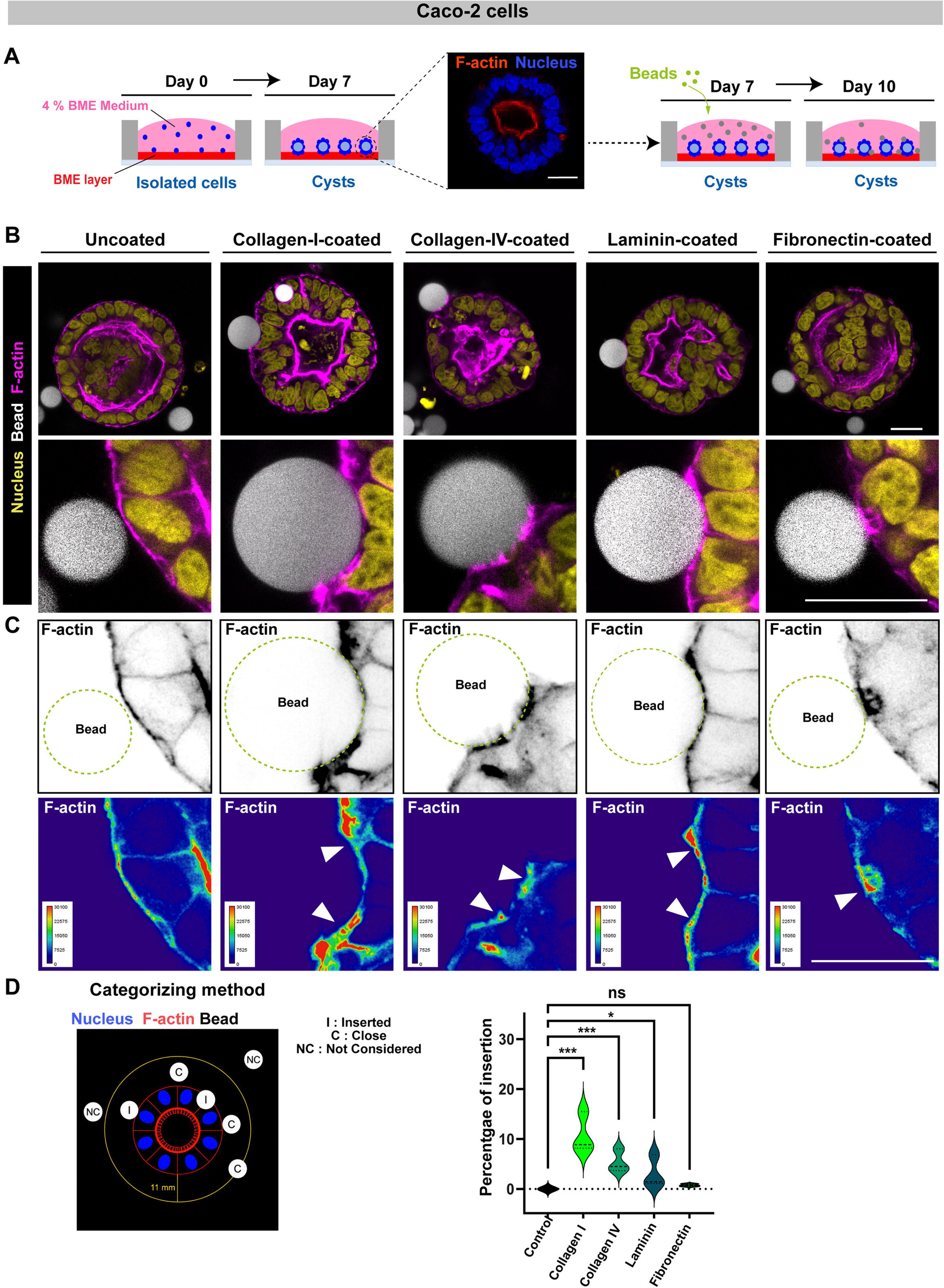
ECM protein functionalization controls tissue insertion efficiency. **A**, Schematic illustrating the procedure for inserting PAAm beads into polarized Caco-2 cysts. Isolated cells were seeded on top of basement membrane extract (BME) and cultured for 7 days in medium supplemented with 4% BME. PAAm beads were then added to the culture medium, and samples were fixed 3 days later. **B,** Confocal analysis of Caco-2 cysts containing uncoated beads (control) or beads coated with collagen-I, collagen-IV, laminin, or fibronectin. Cysts were labelled with phalloidin (magenta) and DAPI (yellow), while PAAm beads are shown in gray. Bottom panels show magnified views of the regions indicated by dashed boxes in the corresponding top panel. Scale bar: 20µm. **C**, Grayscale images show the F-actin channel alongside the corresponding actin intensity heatmaps. Arrowheads indicate regions of F-actin enrichment at the bead-cell interface. The green dashed line represents the position of the beads. Scale bar, 10µm. **D,** Quantification of PAAm bead insertion efficiency. The left panel illustrates the quantification method. The middle panel shows box plots representing statistical analysis of bead insertion across different ECM coatings. n (control uncoated beads) = 290, n (collagen-I-coated beads) =219, n (collagen-IV-coated beads) = 502, n (laminin-coated beads) = 265, n (fibronectin-coated beads) =323. Control vs collagen-I p-value= 1.299e-09; control vs collagen-IV p-value= 3.675e-05; control vs laminin p-value= 0.01619; control vs fibronectin p-value= 0.2866; collagen-I vs collagen-IV p-value= 0.004654; collagen-I vs laminin p-value= 3.917e-05; collagen-I vs fibronectin p-value=1.656e-08; collagen-IV vs laminin p-value=0.04887; collagen-IV vs fibronectin p-value= 0.0004303; laminin vs fibronectin p-value=0.2014.

Polarized epithelial 3D cysts were generated by culturing Caco-2 cells on soft basement membrane extract (BME) for 7 days, forming well-defined monolayers enclosing a central lumen. After cyst polarization (day 7), PAAm beads were added to the culture medium and incubated for 72h until day 10 (Figure 2A). To determine the optimal functionalization for PAAm bead integration, we tested a range of ECM-mimetic coatings, including collagen-I, collagen-IV, laminin, and fibronectin. Importantly, bead insertion did not compromise epithelial architecture: cyst polarity, lumen integrity, and monolayer organization remained intact across all conditions (Figure 2B, Supplementary Figure 2A). Fluorescent labelling revealed that bead–cyst interactions were strictly dependent on bead surface biochemistry: uncoated beads failed to adhere, while ECM-coated beads exhibited coating-specific behaviours, ranging from surface attachment to partial or complete engulfment (Figure 2B-C). Quantitative analysis showed that collagen-I coatings achieved the highest insertion frequency (12.2 ± 4.68%), followed by collagen-IV, laminin, and fibronectin (Figure 2D). Additionally, collagen-I promoted the deepest bead integration within the epithelial layer (Figure 2B-C). In all cases, bead integration was accompanied by localized actin recruitment (Figure 2C), indicating an adaptive cellular response at the bead-tissue interface.

These results demonstrated that ECM surface biochemistry is a critical design parameter for regulating the mechanical coupling between hydrogel microbeads and epithelial tissues. The pronounced efficacy of collagen-I aligned with prior 2D studies, which showed enhanced adhesion and FA signalling of Caco-2 cells on collagen-I substrates [43]. This consistency reinforced the translational relevance of our biohybrid bead interface.

### Collagen-I–functionalized beads elicit tension-bearing focal adhesions

Given its superior insertion efficiency, collagen-I was selected for subsequent mechanistic analyses. Cells surrounding inserted beads exhibited pronounced local shape remodelling, characterized by basal surface curvature directed toward the bead (Figures 2B, 3A), while distal cyst regions retained canonical epithelial morphology (Figure 2B, Supplementary Figure 2A). These observations indicated a spatially restricted mechanical adaptation to bead insertion.

To resolve adhesion mechanics at the bead–cell interface, we examined FA components alongside the actin cytoskeleton. FAs positive for paxillin and its active form, i.e. phospho-paxillin (pPaxillin), were detected at both the basal epithelial surface and at the bead interface, where they aligned with the end of actin bundles (Figure 3A, Supplementary Figure 2B). This organization was characteristic of FAs that transmit forces and anchor actin stress fibers.

**Figure 3:**
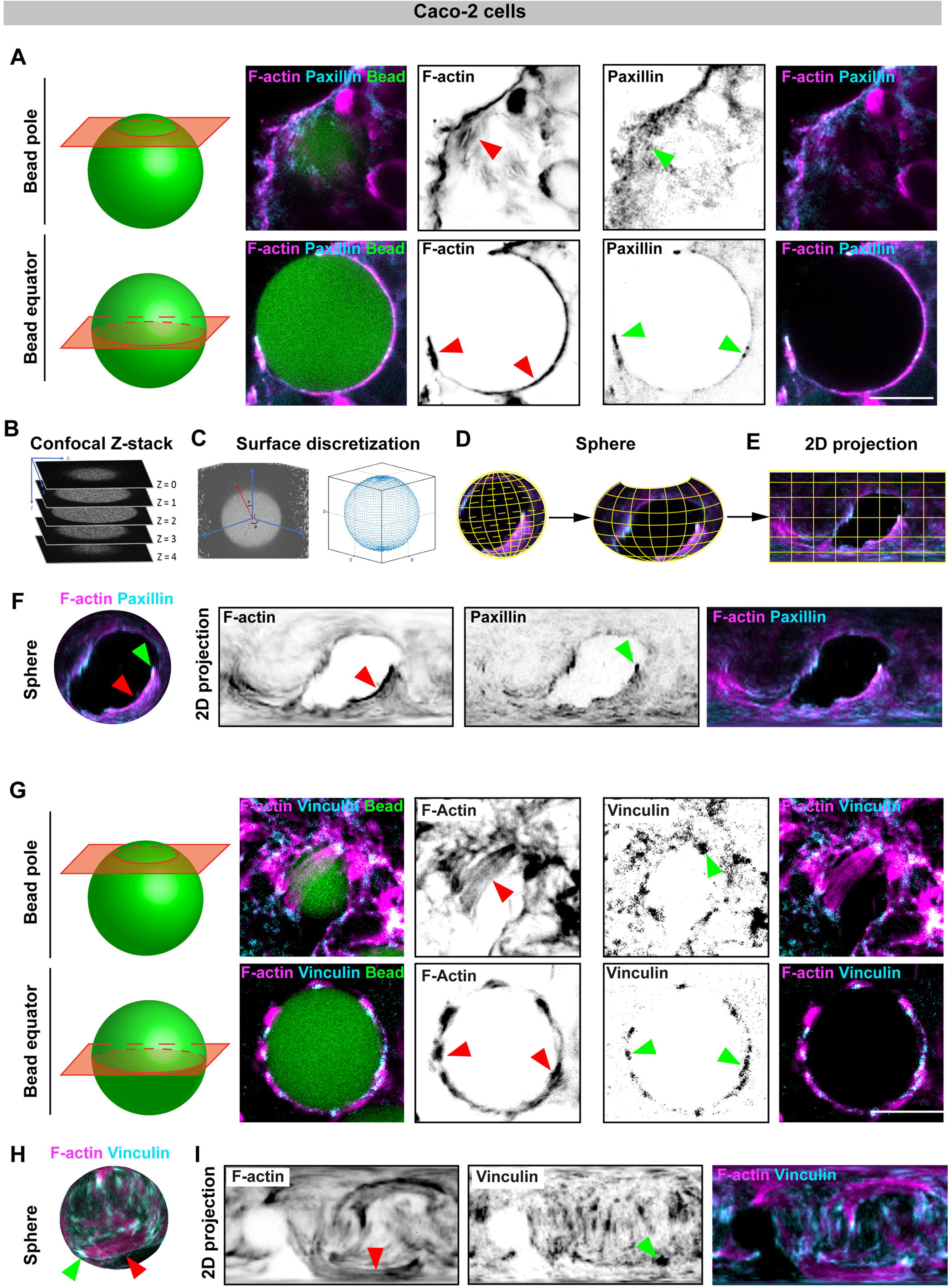
Paxillin and vinculin immunostaining show tension-bearing focal adhesions at the interface between epithelial cells and collagen-I-coated PAAm beads. **A**, Left, schematic representation defining the pole and equatorial imaging planes of the PAAm beads. Right, confocal analysis of the pole and equatorial planes of a collagen-I–coated bead interacting with Caco-2 cyst cells (F-actin: magenta; paxillin: cyan; bead: green). Grayscale images show individual fluorescence channels, while merged images illustrate their combined spatial organization. Red and green arrowheads indicate regions enriched in F-actin and paxillin, respectively, at the bead-cell interface. Scale bar, 10µm. **B-E**, Workflow for 3D reconstruction, including z-stack image acquisition (**B**), coordinate transformation (**C**), projection onto a spherical reference frame (**D**) and projection into a 2D representation of F-actin and paxillin organization on the bead-cell interface (**E**). **F**, Resulting 3D spherical mapping and 2D projection of paxillin (cyan) and F-actin (magenta) organization at the bead-cell interface. Red and green arrowheads indicate regions enriched in F-actin and paxillin, respectively, at the bead-cell interface. **G**, Left, schematic representation defining the pole and equatorial imaging planes of the beads. Right, confocal analysis of the pole and equatorial planes of a collagen-I–coated bead interacting with cells (F-actin: magenta; vinculin: cyan; bead: green). Grayscale images display individual fluorescence channels, and merged images illustrate their spatial organization. Arrowheads indicate regions of F-actin (red) and vinculin (green) enrichment at the bead-cell interface. Scale bar, 10µm. **H**, 3D spherical mapping and 2D projection representation of vinculin (cyan), F-actin (magenta), and their merged visualization at the bead**-**cell interface. Arrowheads indicate regions of F-actin (red) and vinculin (green) enrichment at the bead-cell interface.

To achieve comprehensive spatial resolution, 3D reconstructions of bead–cell interfaces were generated from confocal z-stack datasets (Figure 3B-C). These reconstructions were then transformed into 2D projections, mapping the entire bead surfaces onto a 2D coordinate system (Figure 3D-F). This approach revealed a dense actin network enveloping the bead, with paxillin concentrated at actin filament endpoints and within regions of FA clustering. Notably, vinculin and zyxin—hallmarks of mechanically engaged FAs [44,45]—were robustly recruited to the bead-cell interface and colocalized with actin bundle ends (Figures 3 G-I, 5A top panel). Prominent vinculin-enriched plaques emerged at sites of multiple actin cable convergence, consistent with zones of elevated tensile load. Together, these findings demonstrated that bead insertion is an active, force-mediated process, driven by cytoskeletal remodelling and stabilized by tension-bearing FAs.

### Actomyosin contractility correlates with bead deformation and force anisotropy

To directly link cytoskeletal activity to force generation, we analyzed actomyosin contractility by assessing phospho-myosin light chain-2 (pMLC). While basal pMLC signal was detectable throughout the epithelium, a marked enrichment was observed at bead–cell interfaces, particularly at protrusive contact sites (Figure 4A). 3D reconstructions, combined with 2D projections, further revealed a preferential localization of pMLC at the leading edges of bead-engulfing cells, where it colocalized with actin-rich protrusions (Figure 4B-C).

**Figure 4:**
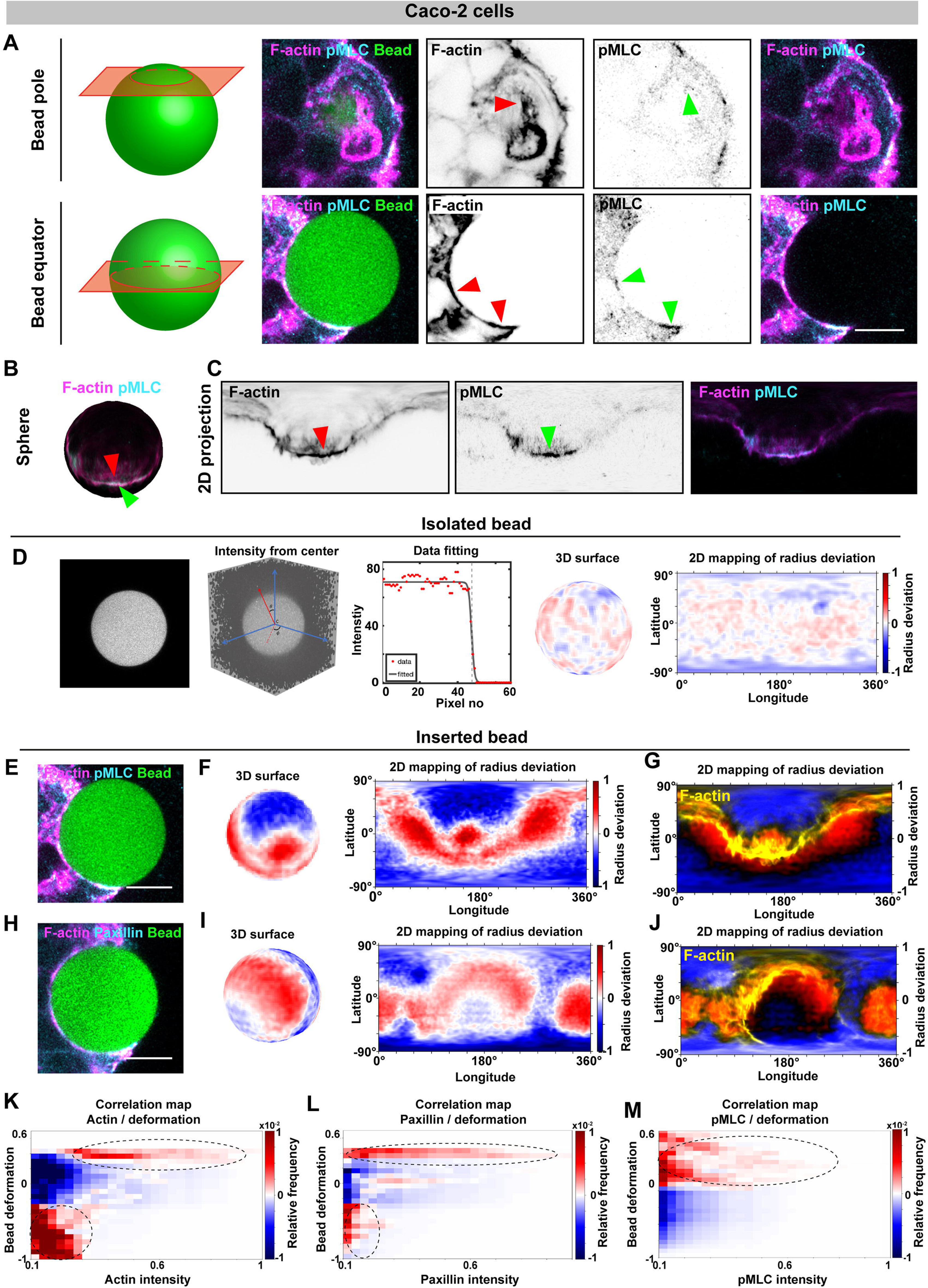
Active actomyosin cytoskeleton at the bead surface correlates with force mapping. **A**, Schematic defining the pole and equatorial imaging planes of the beads. Confocal analysis at the pole and equatorial planes of a collagen-I–coated bead interacting with cells (F-actin: magenta; pMLC: cyan; bead: green). Grayscale images display individual fluorescence channels, and merged images illustrate their spatial organization. Arrowheads indicate regions of F-actin (red) and pMLC (green) enrichment or depletion at the bead-cell interface. Scale bar: 10 µm. **B-C**, 3D spherical mapping (**B**) and 2D projection representation (**C**) of pMLC (cyan), F-actin (magenta), and their merged visualization at the bead-cell interface. Arrowheads indicate regions of F-actin (red) and pMLC (green) enrichment at the bead-cell interface. **D**, Workflow for computing bead deformation, including confocal acquisition of an isolated bead, coordinate transformation, data fitting to an ideal sphere (data: red dots; fit: black line), projection onto a spherical reference frame, and conversion into a 2D representation. Positive radius deviations as compared to the average bead radius are shown in red, and negative radius deviations in blue. **E, H,** Collagen-I-coated bead inserted into a Caco-2-cyst, stained for F-actin (magenta) and pMLC (cyan, **E**) or paxillin (cyan, **H**). **F, I**, 3D reconstruction of the deformed bead surface and its 2D projection representation. Positive radius deviations are shown in red, and negative radius deviations in blue. **G, J**, Overlay of the 2D projection representation and actin signal (yellow). **K-M**, Representative correlation maps of the deformations (left, y-axis), the actin intensity (**K**, x-axis), paxillin intensity (**L**, x-axis), pMLC intensity (**M**, x-axis) and frequency of occurrence (right, y-axis). High frequency is indicated in red, and low frequency in blue.

To determine whether localized contractility translated into measurable mechanical outputs, bead deformations were quantified by calculating radial deviations from an ideal sphere (Figure 4D). Isolated beads exhibited minimal and spatially uniform deviations (0.09 ± 0.04µm), reflecting fabrication-associated baseline heterogeneity (Figure 4D, Supplementary Figure 1D-E). In contrast, cyst-inserted beads displayed pronounced spatial heterogeneity in their deformation patterns, with localized indentations and expansions averaging at 0.21 ± 0.09µm, indicative of anisotropic cellular force application (Figure 4E-J). Thus, integration of molecular and mechanical datasets allowed the direct mapping of cytoskeletal organization onto deformation patterns. This analysis revealed a clear correlation between actin arrangement and force modality: regions denser in actin correlated with stronger pulling forces as compared to regions that are more dilute in actin (Figure 4F-G, I-J). To build on these observations, we performed a correlation analysis at the bead-cell interface (Figure 4K-M). Regions of high actin intensity, corresponding to dense protrusive structures, exhibited an increased probability of positive radius deviation, suggesting more tensile (i.e. pulling) forces. In contrast, regions of lower actin intensity were preferentially associated with more compressive (i.e. pushing) forces (Figure 4 K). Consistently, paxillin-enriched protrusive domains also correlated primarily with more tensile deformation signatures (Figure 4L). In agreement with numerous 2D studies and the broader literature, our correlation analysis demonstrated that pMLC enrichment was preferentially associated with positive radius deviations, consistent with stronger pulling forces (Figure 4M).

To assess the contribution of myosin-II-dependent actomyosin contractility to these spatial force patterns, we inhibited myosin-II activity using blebbistatin. Following treatment, circumferential actin cables appeared largely decoupled from the bead surface, although residual actin structures remained associated with the bead-cell interface (Figure 5A). As expected, pMLC levels were markedly decreased upon blebbistatin treatment (Supplementary Figure 3A-C). Consistent with impaired force transmission at cell-bead adhesion sites, zyxin—a marker of mechanically loaded FAs—was also reduced (Figure 5A-C). Notably, bead deformation was diminished following blebbistatin treatment (Figure 5D-G, Supplementary Figure D-G), demonstrating that actomyosin contractility is a primary driver of the forces exerted on the bead surface. The correlation analysis was therefore performed under this condition. Blebbistatin markedly reorganized the relationship between local actin intensity and bead deformation. In contrast to control conditions (i.e. without blebbistatin), where local actin intensity correlated with local bead deformation, blebbistatin treatment disrupted this actin–bead deformation correlation, making it significantly weaker (Figure 5H-J, Supplementary Figure 3H-J). Thus, inhibition of myosin-II activity disrupted the characteristic coupling between local actin organization and bead deformation, suggesting that actomyosin contractility is required to establish or maintain the spatial organization of distinct pulling and pushing force regimes at the cell–matrix interface.

**Figure 5:**
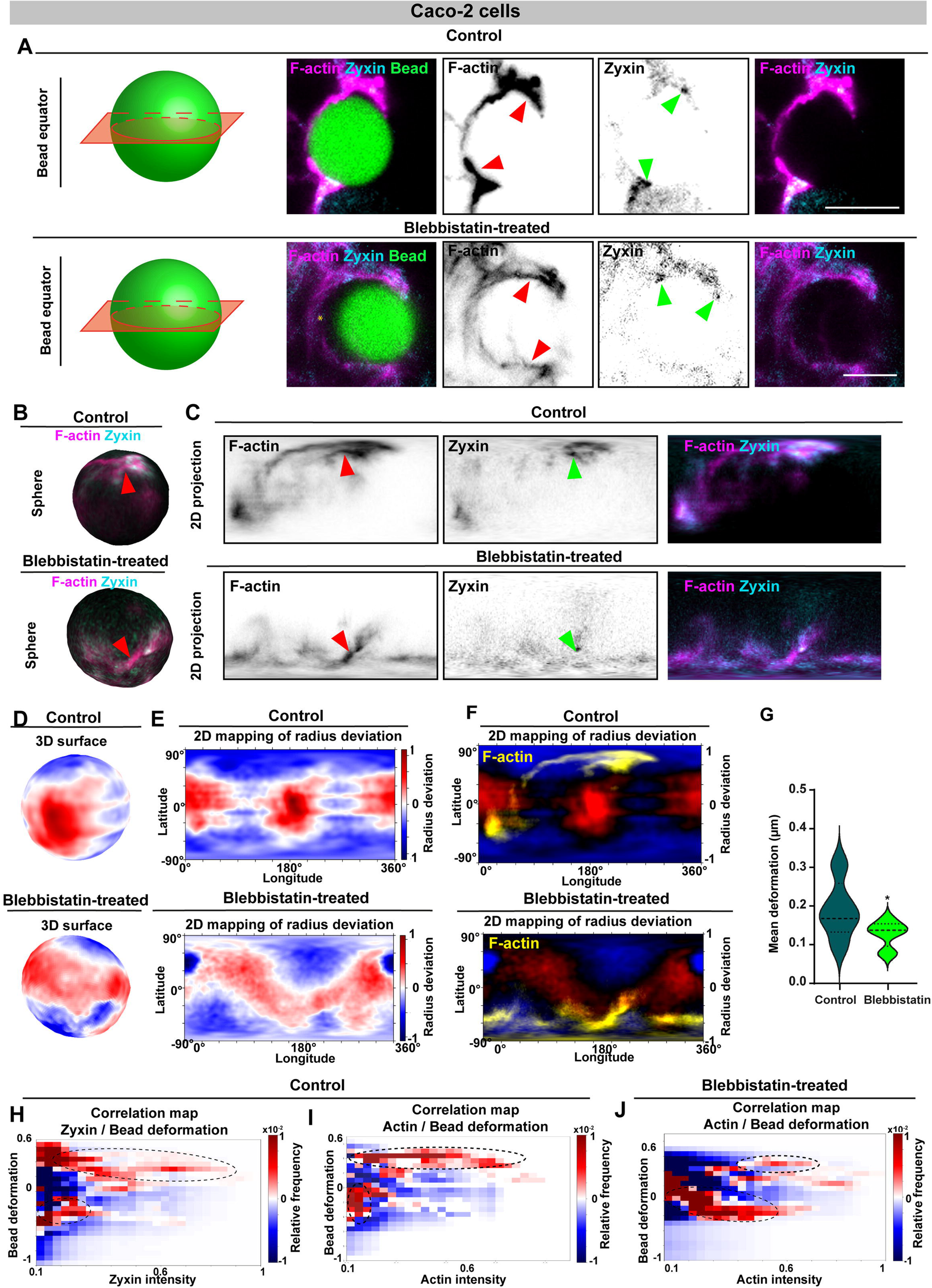
Contractility inhibition decouples bead deformation from actin adhesion at the bead surface. **A**, Schematic defining the equatorial imaging planes of the beads. Confocal analysis at the equatorial planes of a collagen-I–coated bead interacting with control cells (top panel) and blebbistatin-treated cells (bottom panel) (F-actin: magenta; zyxin: cyan; bead: green). Grayscale images display individual fluorescence channels, and merged images illustrate their spatial organization. Arrowheads indicate regions of F-actin (red) and zyxin (green) enrichment or depletion at the bead-cell interface. Scale bar: 10 µm. **B-C**, 3D spherical mapping (**B**) and 2D projection representation (**C**) of zyxin (cyan), F-actin (magenta), and their merged visualization at the bead-cell interface for control (top panel) and blebbistatin-treated cells (bottom panel). Arrowheads indicate regions of F-actin (red) and zyxin (green) enrichment at the bead-cell interface. **D**,-**E,** 3D reconstruction of the deformed surface of the bead (**D**) and 2D projection representation (**E**) for control (top panel) and blebbistatin-treated (bottom panel) cells. Positive radius deviations are shown in red, and negative radius deviations in blue. **F**, Overlay of the 2D projection representation and actin signal (yellow) for control (top panel) and blebbistatin-treated (bottom panel) cells. **G,** Quantification of PAAm bead deformation in blebbistatin-treated (black) and control (green) cells. Violin plots represent statistical analysis of bead deformation (in µm) under control and blebbistatin-treated conditions. N= 2 for control and blebbistatin-treated samples, n = 5 control beads, n= 9 blebbistatin-treated beads. Mann-Whitney test, *p < 0,05. **H-J,** Representative correlation maps of bead deformations (y-axis left), zyxin intensity in control conditions (**H**, x-axis), actin intensity in control conditions (**I**, x-axis), actin intensity in blebbistatin-treated conditions (**J**, x-axis) and frequency of occurrence (right, y-axis). High frequency is shown in red, and low frequency in blue.

Altogether, these findings revealed a mechanical duality in which epithelial cells exert spatially patterned, anisotropic forces. PAAm microbeads therefore emerge as embedded force sensors within epithelial tissues, enabling quantitative analyses of how localized actomyosin contractility generates patterns of coordinated interaction forces that sculpt and deform the 3D microenvironment.

### Functionalization of PAAm beads with adherens junction components

We next investigated how functionalization with E-cadherin, a central mediator of cell-cell adhesion [46], modulated bead–cell interactions within polarized 3D epithelia. Control IgG-Fc-coated beads failed to integrate into the cysts because they did not attach to the epithelial cells (not shown). Exposure to E-cadherin-Fc-coated beads did not perturb the overall epithelial architecture (Figure 6A), indicating that tissue integrity was preserved following E-cadherin-coated bead insertion, as observed for ECM-based functionalization. E-cadherin-coated bead integration induced a significant recruitment of actin (Figure 6A-B) and β-catenin (Figure 6C) at the bead-cell interface, reflecting an adaptive cell response to the adhesion cue. Thus, E-cadherin bead functionalization was sufficient to mimic native cell-cell interaction and promote the assembly of canonical AJ-like structures at the bead surface.

**Figure 6:**
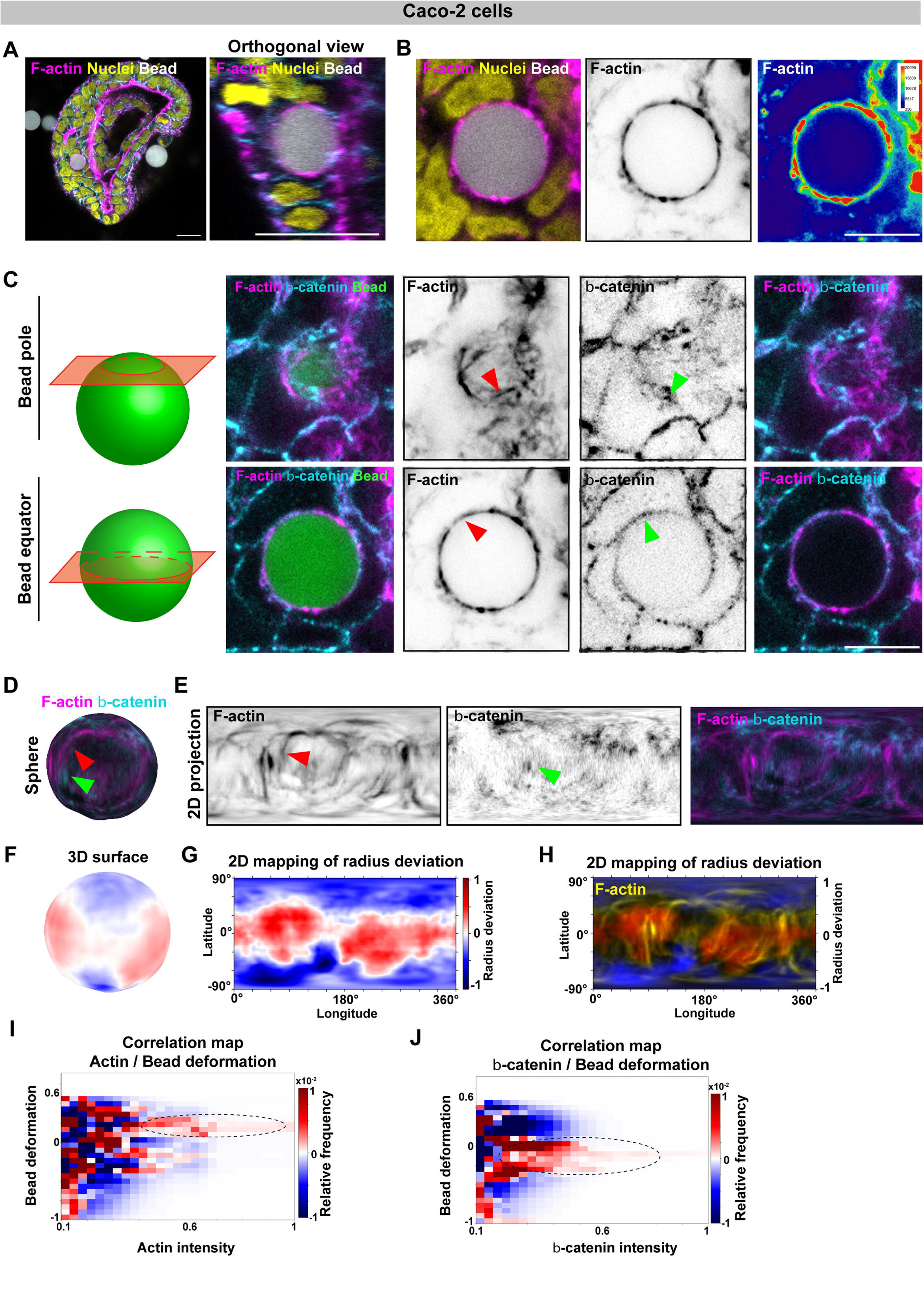
E-cadherin protein functionalization correlates with force mapping. **A,** Confocal analysis of Caco-2 cysts with E-cadherin-coated beads, labelled with phalloidin (magenta), DAPI (yellow) and beads (gray). Left panel shows a zoomed zy-view of the bead. Scale bar: 20µm. **B**, Confocal analysis of actin at the bead surface (actin: magenta; bead: gray; nuclei: yellow). Grayscale image shows the actin channel with its heatmap representation on the left. Scale bar: 10µm. **C**, Left, schematic defining the pole and equatorial imaging planes of the beads. Right, confocal analysis at the pole and equatorial planes of a collagen-I–coated bead interacting with cells (F-actin: magenta; β-catenin: cyan: bead: green). Grayscale images display individual fluorescence channels, and merged images illustrate their spatial organization. Arrowheads indicate regions of F-actin (red) and β-catenin (green) enrichment or depletion at the bead-cell interface. Scale bar: 10 µm. **D-E,** 3D spherical mapping (**D**) and 2D projection representation (**E**) of β-catenin (cyan), F-actin (magenta), and their merged visualization at the bead-cell interface. Arrowheads indicate regions of enrichment of F-actin (red) and β-catenin (green) at the bead-cell interface. **F-G,** 3D reconstruction of bead surface deformations (**F**) and 2D projection representation (**G**). Positive radius deviations are shown in red, and negative radius deviations in blue. **H,** Overlay of the 2D projection representation and actin signal (yellow). **I-J**, Correlation maps of the deformation (left, y-axis), the actin intensity (**I**, x-axis), β-catenin intensity (**J**, x-axis) and frequency of occurrence (right, y-axis). High frequency is shown in red, and low frequency in blue.

3D reconstructions and 2D projections further revealed that β-catenin was distributed circumferentially around the bead and colocalized with an organized actin network (Figure 6D-E). This spatial organization was consistent with the formation of a continuous junctional complex encasing the bead, analogous to endogenous AJs in epithelial tissues. However, in contrast to ECM-coated beads, no prominent actin bundles were detected at the surface of E-cadherin-coated beads. Instead, a thin actin network partly colocalized with β-catenin was observed (Figure 6D-E). Moreover, as observed for ECM-coated beads, insertion of E-cadherin-coated beads led to pronounced spatial heterogeneity in bead deformation patterns (Figure 6F-H).

Correlation analysis revealed no clear relationship between actin organization at E-cadherin-mediated junctions and bead deformation patterns (Figure 6I,J). Regions of high actin intensity were slightly associated with regions of somewhat positive radius deviation (Figure 6I), while β-catenin intensity was slightly associated with regions of somewhat negative radius deviation (Figure 6J). This spatial association suggested that cell–cell adhesion sites enriched in β-catenin were preferentially associated with bead deformations consistent with a more compressive–or at least less tensile forces–at the bead interface. Because β-catenin was specifically recruited to E-cadherin–mediated adhesion sites, these results further suggested that the anchoring points established at the bead surface sustain more compressive–or at least less tensile–forces than other regions. Strikingly, this behaviour contrasted with that observed at ECM-mediated contact sites, where paxillin-enriched structures were more clearly linked to tensile forces. Together, these findings revealed a fundamental difference in how epithelial cells may mechanically engage cell-cell versus cell-matrix adhesions in 3D conditions.

### Translation to intestinal organoid systems

To assess the broader applicability of the PAAm microbead-based force-sensing platform in physiologically relevant settings, we extended our approach to primary mouse intestinal organoids (Figure 7A). Unlike transformed Caco-2 cells, intestinal organoids recapitulate key features of native intestinal architecture, including crypt- and villus-like domains (Figure 7A), while exhibiting a softer mechanical microenvironment characteristic of healthy intestinal tissue [47]. Accordingly, we employed soft PAAm beads (∼600Pa) to approximate the elastic modulus of healthy intestinal tissue, and stiff PAAm beads (∼10kPa) to model the increased stiffness associated with intestinal adenocarcinoma (Figure 7B,I). Collagen-I-functionalized beads were introduced into dissociated organoid cultures using the same non-invasive incubation strategy employed for Caco-2 cysts, enabling spontaneous bead incorporation without mechanical disruption (Figure 7A-B). After four days in culture, collagen-I-coated beads were successfully incorporated into the crypt domain of organoids under both stiffness conditions (Figure 7B, D, I, K; Supplementary Figure 4A-B). Importantly, bead insertion did not disrupt organoid epithelial architecture. Apico-basal polarity, lumen integrity, and the organization of the epithelial monolayer into crypt and villus domains were preserved in both soft- and stiff-bead conditions (Figure 7B, I). In contrast, actin organization at the bead-tissue interface differed between the two conditions (Figure 7C, J). Whereas soft beads were associated with a relatively homogeneous actin distribution, stiff beads induced a pronounced reorganization of the actin cytoskeleton at the bead-cell interface (Figure 7C red arrowheads), indicative of an adaptative cellular response to a mechanically stiffer microenvironment. To achieve comprehensive spatial resolution, we applied the established pipeline for 3D reconstructions of the bead–cell interface followed by 2D surface mapping. This analysis revealed marked differences in actin organization at the bead-cell interface between stiff and soft conditions (Figure 7D, E and K,L) whereas pronounced bead deformations were observed in both cases (Figure 7F, M). Organoids containing embedded beads displayed spatially heterogeneous bead deformation patterns, indicative of anisotropic force generation by epithelial cells. Notably, these deformations were greater than those measured in Caco-2 cysts, indicating that intestinal organoids generated larger mechanical bead deformations in their 3D microenvironment. Furthermore, stiff beads in contact with crypt domain of organoids are significative compressed with no apparent tensile deformation (Figure 7F), whereas soft beads exhibited both compressive and tensile deformation (Figure 7M).

**Figure 7:**
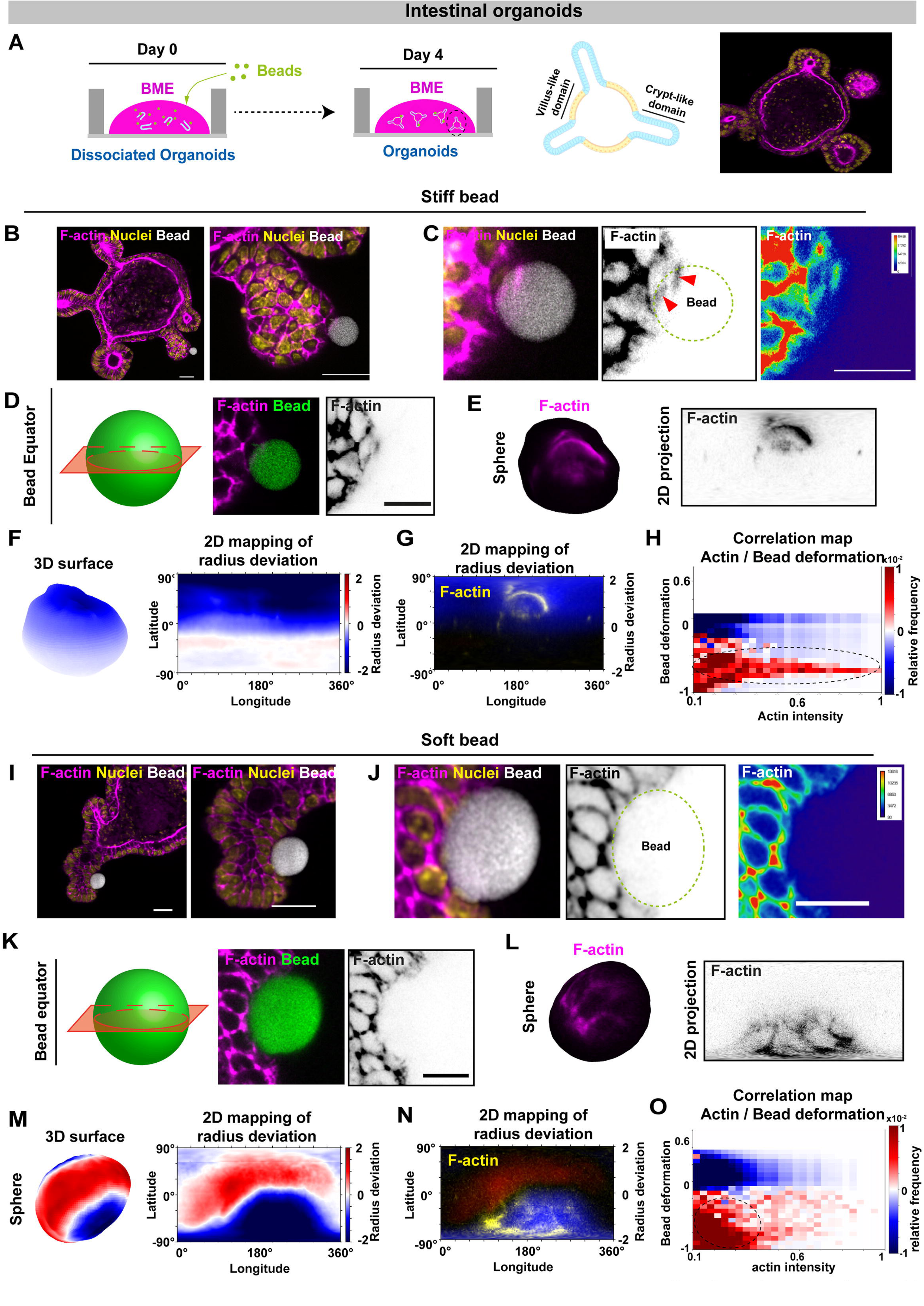
Application of the approach to the intestinal organoid model. **A**, Schematic of the procedure to grow organoids with beads. Dissociated organoids are mixed with beads and embedded in BME. After 4 days in culture, differentiated organoids form, containing villus-like and crypt-like domains. Left panel shows an organoid labelled for F-actin (magenta) and DAPI (yellow). Stiff (10kPa) beads (**B-H**) and soft (600Pa) beads (**I-O**) are engulfed by the organoids. For each condition, beads are inserted into the crypt domain (**B, I**). Organoids are labelled for F-actin (magenta), nuclei (DAPI: yellow). Scale bar 20µm. **C, J,** Confocal analysis of actin at the surface of stiff (**C**) or soft (**J**) bead (actin: magenta; bead: gray; nuclei: yellow). Grayscale images show the actin channel with its heatmap representation on the left. Scale bar: 10µm. **D, K,** Left, schematic defining the equatorial pole imaging planes of the beads. Right, confocal analysis at the equatorial plane of a collagen-I–coated stiff (**D**) or soft (**K**) bead interacting with cells (F-actin: magenta; bead: green). Grayscale images display individual fluorescence channels, and merged images illustrate their spatial organization. Scale bar 10 µm. **E, L**, 3D spherical mapping (left) and 2D projection representation (right) of F-actin (magenta and gray, respectively), when insertion of stiff (**E**) or soft (**L**) bead. **F, M**, 3D reconstruction of the deformed bead surface (left) and 2D projection representation (right), when insertion of stiff (**F**) or soft (**M**) bead. Positive radius deviations are shown in red, and negative radius deviations in blue. **G, N**, Overlay of the 2D projection representation and actin signal (yellow), when insertion of stiff (**G**) or soft (**N**) bead. **H, O,** Representative correlation maps of the bead deformation (left, y-axis), actin intensity (x-axis), and frequency of occurrence (right, y-axis), when insertion of stiff (**H**) or soft (**O**) bead. High frequency is shown in red, and low frequency in blue.

To further investigate these observations, we performed a correlation analysis at the bead-cell interface (Figure 7H,O). Bead stiffness modulated the relationship between actin organization and mechanical bead deformation within the crypt domain. On stiff beads, intermediate to high actin intensities were associated with compressive deformation, consistent with pushing regime (Figure 7H). These data suggested that actin structures were efficiently coupled to force transmission under stiff conditions in the crypt domain. In contrast, on soft beads, the relationship between actin intensity and deformation was limited to low actin intensities (Figure 7O), suggesting less defined associated actin structures. The distribution lacked a well-defined structure, with a broad range of deformation values that were not strongly associated with high actin intensity levels (Figure 7O). Although tensile deformations and pulling forces were still observed, their association with actin organization was more diffuse (Figure 7O), indicating a different coupling between cytoskeletal architecture and mechanical output. Together, these findings suggested that, in a softer mechanical environment, forces generated by the actin network were more broadly distributed and transmitted to the surrounding matrix.

## Discussion

Cells sense and respond to mechanical forces through continuous remodelling of the actin cytoskeleton, cell–matrix and cell-cell adhesions, a process extensively characterized in 2D systems but still incompletely understood in 3D environments, where geometry, confinement, and epithelial polarity introduce additional layers of complexity [22,48]. In this work, we leverage recent advances in 3D elastic polyacrylamide (PAAm) bead–based force sensors [34,36] to locally perturb and quantify mechanical interactions within highly organized Caco-2 epithelial cysts and mouse intestinal organoids. To our knowledge, this represents the first implementation of deformable PAAm bead force sensing in polarized 3D epithelial models. We developed and validated a versatile PAAm microbead platform that enables both mechanical manipulation and force readout at the single-cell scale within 3D cultures. Our results demonstrate that these beads can be engineered with tunable stiffness, controlled size, and stable protein functionalization, making them robust mechanical probes in physiologically relevant epithelial environments.

The bead fabrication strategy provides access to a wide range of physiologically relevant elastic moduli, spanning those reported for healthy to pathological intestinal tissues (∼600 Pa to ∼7.5 kPa). This tunability is particularly valuable for modelling mechanical transitions associated with disease progression, such as tumorigenesis. By controlling stiffness through the acrylamide/bis-acrylamide ratio and selecting bead size via sequential filtration, we produced cell-scale beads that closely replicate the biomechanical environment of the intestinal epithelium. The beads are purely elastic, allowing for long-term observations without the measurements being affected by viscous relaxation. Additionally, homogeneous fluorescent labelling of the entire bead volume allowed precise 3D shape reconstruction and reliable quantification of deformation. This enabled dense, continuous mapping of both compressive and tensile deformations across the bead surface.

Unlike probes with discretely labels that rely on interpolation between sparse fiducial markers, our approach delivered a dense, continuous deformation field across the bead surface. This enhanced spatial resolution and sensitivity, particularly under complex or asymmetric loading conditions. Compared to other force-sensing strategies, PAAm beads offered several key advantages. Unlike oil microdroplets, which are fluid and isotropic and thus limit force readouts to surface tension– dominated deformations, PAAm beads behaved as elastic solids with well-defined, tunable mechanical properties. This allowed them to capture both compressive and tensile deformations while better mimicking the mechanics of soft tissues. As localized mechanical transducers, PAAm beads simplified analysis, reduced computational demands, and enabled direct correlation of force readouts with specific cellular or subcellular structures. Together, these features established PAAm beads as a powerful, adaptable tool for probing epithelial mechanics in 3D.

Our findings also demonstrated that ECM composition critically governs bead–cyst interactions, highlighting the interplay between biochemical and mechanical cues in mechanotransduction. Collagen-I–coated beads showed the highest insertion efficiency, consistent with prior 2D studies that report preferential adhesion of Caco-2 cells to collagen-I via integrin-mediated signalling pathways [49,50]. This behaviour likely reflects the expression of collagen-I–binding integrins, such as α2β1, and their capacity to form mechanically robust FAs. In contrast, coatings with collagen-IV, laminin, or fibronectin resulted in weaker interactions, emphasizing the specificity of ECM in adhesion dynamics. High-resolution confocal imaging revealed that collagen-I–coated beads induced the formation of mature FAs, marked by the enrichment of paxillin and vinculin at the ends of actin filaments. The colocalization of vinculin, paxillin and zyxin with converging actin stress fibers indicated load-bearing adhesions, consistent with tension-transmitting FAs described in 2D systems [51,52]. Phosphorylated myosin light chain-2 (pMLC) was enriched at bead–cell interfaces, particularly at protrusive leading edges engulfing the ECM-coated beads, linking actomyosin contractility to measurable bead deformations. This supported the notion that bead insertion is an active, force-driven process mediated by actomyosin contractility. Quantitative probabilistic correlation analyses further revealed that actin-dense protrusions and paxillin-positive puncta were strongly associated with tensile deformations indicating pulling forces, whereas extended circumferential actin cables corresponded to compressive (i.e. pushing) deformations. These results suggested a dual mechanical strategy in which epithelial cells both pull on and push against the bead to remodel their local microenvironment during insertion. This coordinated pushing–pulling mechanism resembles adaptive force strategies recently described in confined cell migration, where cells generate pushing forces at the rear and pulling forces at the front to overcome spatial constraints [53].

E-cadherin-coated beads, which mimic cell–cell adhesion cues, integrated efficiently without disrupting epithelial polarity or lumen organization. They induced the recruitment of β-catenin and a thin circumferential actin network at the bead interface. Importantly, bead integration did not disrupt overall tissue architecture, demonstrating that localized mechanical probing can be performed without compromising global epithelial organization. E-cadherin-functionalized beads further revealed how adhesion context shapes force generation. Unlike ECM-coated beads, E-cadherin-coated beads recruited thin, circumferential actin networks along with β-catenin at the bead surface, resembling AJ-like assemblies. Probabilistic correlation analyses showed that β-catenin–enriched regions primarily generated compressive deformations, compatible with pushing forces, while actin intensity exhibited heterogeneous associations with both pulling and pushing regimes. This underscored a fundamental distinction between cell–cell and cell–matrix interactions in 3D: ECM contacts may favour actomyosin-mediated pulling, whereas AJs may drive compressive forces, indicating adhesion-specific mechanical strategies.

Extending these observations to mouse intestinal organoids demonstrated the broader applicability of the approach. Beads inserted into organoid crypt domains displayed highly anisotropic deformations– greater than those in Caco-2 cysts–reflecting the higher force-generating capacity of organoid epithelia. Bead stiffness also modulated the coupling between actin organization and mechanical output: in stiff microenvironments, actin intensity strongly correlated with compressive deformation compatible with pushing forces, whereas in soft environments, this relationship was weaker and more diffuse. These results suggested that tissue stiffness critically influences the efficiency of cytoskeletal force transmission in 3D, with stiff environments enhancing directional coupling of actin to mechanical output.

Collectively, these findings established a robust materials-based framework for probing mechanical interactions in 3D epithelial systems, bridging molecular-scale force mapping with tissue-level organization. Beyond force readout, this platform can be extended to investigate the effects of ECM composition, stiffness gradients, or pharmacological perturbations on cellular force dynamics in complex 3D contexts, such as organoids, tumours, and regenerative models. Coupling PAAm bead– based force sensing with live imaging or advanced traction reconstruction approaches promises unprecedented insights into how epithelial cells sense, interpret, and remodel their mechanical environment during development, disease progression, and cancer invasion.

## Acknowledgments

This work was supported by doctoral fellowships to N.G and M.I from Turing Centre for Living Systems, and by grants from the CNRS through the MiTi interdisciplinary programs (to E.B and M.M, to D.D.), the French Foundation ARC (to D.D), the Agence Nationale de la Recherche (ANR-25-CE13-3505-01 to D.D., ANR-16-CONV0001 to M.M), and the 2021-2023 Cancer Control strategy, on funds administered by INSERM (to D.D.), France 2030, the French and from the Excellence Initiative of Aix-Marseille University–A*MIDEX, the European Research Council (ERC) under the Marie Curie Sklodowska action (MSCA-IF, to C.V.), the ITMO Cancer of Aviesan on funds Cancer-2021 (ATIP Avenir to C.V.). E.B, D.D, M.M, A.L-B., C.V are CNRS staff members, D.M.H is INSERM staff and F.R. is Professor at Aix Marseille University. We thank the imaging facility at IBDM, member of the National Infrastructure France-BioImaging (https://ror.org/01y7vt929) supported by the French National Research Agency (ANR24-INBS-0005 FBI BIOGEN).

## Supplemental Information

Supplementary Figures S1 to S2.

## Author contributions

N.G, M.I, B.L., C.V, D.M.H., D.D., M.M., and E.B. designed and performed experiments and required analyses. N.G, M.I, A.L.B., F.R., D.D., M.M., and E.B. coordinated the overall research and experiments, and wrote the manuscript.

## Material and methods

### Polyacrylamide bead production

Polyacrylamide beads were synthesized via a PBS/oil emulsion method. The oil phase consisted of 8% (w/v) Krytox 157 (FSHDNK157FSH500G Samaro) in HFE-7500 3M (FluoroChem). The aqueous phase contained 186µl acrylamide (BioRad), 80µl bis-acrylamide (BioRad) for 10kPa stiffness or 22µl bis-acrylamide for 800Pa stiffness, 0,00001% of Fluorescein-O-methacrylate solution (Sigma-Aldrich), and PBS to a final volume of 1 ml. 0,004% ammonium persulfate (Sigma) and 0,02% TEMED (Sigma) were added in the aqueous phase mixture to initiate the polymerization. The emulsion was generated using a 25G x5/8” needle. Polymerization was achieved at room temperature for 1 hour. Post-polymerization, beads were washed multiple times with 20% 1H,1H,2H,2H-Perfluoro-1-octanol (#11490701, Fisher Scientific) in HFE 7500 3M. After centrifugation at 4,500xg for 1 min, beads were segregated into the PBS phase. The supernatant was gently aspirated by pipetting, and beads were left to swell overnight at 4°C. Finally, beads were centrifuged again at 4,500xg for 1 min, washed in PBS, filtered (PluriSelect Strain), and counted using a hemocytometer.

### Atomic Force Microscopy (AFM)

Beads were immobilized on a CellTak-coated Petri dish (Corning, DLW354240). AFM measurements were performed using a NanoWizard IV AFM (JPK-Bruker), coupled to a TiU inverted microscope (Nikon) equipped with a 20x objective. MLCT-Bio_D cantilevers (V-shaped cantilevers, four-sided regular pyramidal tips with 35° semi-open angle, nominal spring constant of 0.03 N/m, Bruker) were used. The spring constant was calibrated in liquid using the thermal spectrum and the Sader method [54]. Then, on top of each bead, 50 force curves were acquired at an approach velocity of 2 µm/s, with a setpoint of 0.5-2nN. This resulted in an indentation depth of 0.5-2.5μm), corresponding to 2 to 10% of the bead diameter, and a contact radius of 0.25 to 1.2 µm. These dimensions are comparable to the deformations induced by the cells, indicating that AFM allows to probe the same scales and mechanical properties as the cells. Curves showing a jump-to-contact or a slope inflexion in the indentation region were discarded. The Young modulus was determined by fitting the approach curves up to 2μm indentation using a numerical model. This model considered the total deformation induced by a pyramid tip on a spherical bead (with a Poisson ratio of 0.5) laying on a flat, rigid surface, including both indentation and sphere deformation against the surface.

To verify the incompressibility of the PAAm beads, we measured their lateral deformation while the AFM tip applied a force to the top. For this, we used a tilt-compensated 20×20 µm platform cantilever (Vmicro) with a nominal spring constant of 0.5 N/m. The spring constant was calibrated in liquid using the thermal spectrum and the Sader method [54]. The bead size and lateral deformation were measured in top view, while indentation was derived from the force-distance curves. These results were validated using optical measurements of deformation and indentation with a ‘side viewer’ cantilever holder (JPK). Bead dimensions were analyzed using ImageJ and the Poisson ratio was calculated as the ratio of relative lateral deformation over the relative indentation (n=15 measurements over 10 beads).

To measure the bead elasticity, MLCT-Bio_D cantilevers (V-shaped cantilevers, four-sided regular pyramidal tips with 35° semi-open angle, nominal spring constant of 0.03 N/m, Bruker) were used. The spring constant was calibrated in liquid using the thermal spectrum and the Sader method [54]. Then, on top of each bead, 50 force curves were acquired at an approach velocity of 2 µm/s, with a setpoint of 0.5-2nN. This resulted in an indentation depth of 0.5-2.5μm), corresponding to 2 to 10% of the bead diameter, and a contact radius of 0.25 to 1.2 µm. These dimensions are comparable to the deformations induced by the cells, indicating that AFM allows to probe the same scales and mechanical properties as the cells. Curves showing a jump-to-contact or a slope inflexion in the indentation region were discarded. The Young modulus was determined by fitting the approach curves up to 2μm indentation using a numerical model. This model considered the total deformation induced by a pyramid tip on a spherical bead (with a Poisson ratio of 0.5) laying on a flat, rigid surface, including both indentation and sphere deformation against the surface.

### Surface functionalization and coating

3.10^6^ beads were centrifugated at 5,000xg for 1 min and washed twice with 0.1M HEPES (#15630-056, Gibco). The bead pellet was resuspended in 2.25mg/ml of Sulfo-SANPAH (diluted in 0.1M HEPES) and irradiated for 11 min under UV light (365nm, 20W UV lamp - #UVPA95-0045-05 XX, VWR International). After irradiation, beads were washed with 0.1M HEPES. Functionalized beads were transferred to LowBind tubes (1.10^6^ beads per tube) for coating.

*ECM coating* – Beads were incubated in 0.1 HEPES with the following protein concentrations: collagen-I: 0.2mg/ml, collagen-IV: 0.025mg/ml, laminin: 0.2mg/ml, fibronectin: 0.4mg/ml, control: 0.1M HEPES solution (no protein). Incubation was performed for 5 hours at 4°C, followed by three PBS washes before use.

*E-cadherin coating* – Beads were first coated with 100µg of Protein-G in 0.1 M NaPhosphate buffer (pH 8) for 1hr at 4°C. They were then incubated with 200µg of E-cadherin-Fc or IgG-Fc proteins (control beads) in NaPhosphate buffer for 2 hours at 4°C. After centrifugation, beads were incubated with a crosslinking buffer (25mM DMP, 0.2M triethanolamine, pH 8.2) for 1 hour. Finally, beads were washed three times with PBS before use.

### Epithelial cell and organoid cultures

*Caco-2 cell culture* – Caco-2 human colonic epithelial cells were cultured as described in [55]. Cells were maintained in DMEM Glutamax (#61965-026, Gibco) supplemented with 20% Fetal Bovine Serum and 1% Minimum Essential Medium Non-Essential Amino Acids (#11140-035, Life Technologies). Cells were incubated at 37°C with 5% CO2.

For 3D cyst cultures, 5,000 Caco-2 cells diluted in 4% Basement Membrane Extract, BME) were seeded in 35 mm glass-bottom dishes (#81218-200, Ibidi) pre-coated with 80 μl of Cultrex Reduced Growth Factor BME type 2 PathClear (Cultrex #3533-005-02, #3532-010-02 BIO-TECHNE). Culture medium was replenished every 2 days, and cysts formed after 7 days of culture. Once cysts were established, the culture medium was removed and replaced with 1ml of media containing 500,000 beads. After 45min, the dish was filled with 4% BME-supplemented culture medium. For inhibition of myosin-II activity, cysts with inserted beads were treated with either 10 µM blebbistatin (#B0560, Sigma-Aldrich) or DMSO (vehicle control). Treatment lasted 1 hour, after which samples were processed for immunostaining.

*Mouse intestinal organoid cultures* – Small intestinal crypts were isolated from 6-12-week-old adult male Villin-Tomato mice. VillinCreERT2-tdTomato organoids were generated from mice provided by Danijela Vignjevic (Insitut Curite, Paris) [56] and breed in the animal house facility of the Institut de Biologie du Développement de Marseille (Marseille). Mice were housed in EOPS (Environment without Specific Pathogenic Organisms) environment, and handled in accordance with French regulation for animal care. Animal procedures were conducted in accordance with the guidelines of the French regulation for animal care. No ethical approval was required, as animal procedures were restricted to mouse sacrifice for organ dissection. Following euthanasia by cervical dislocation, the small intestine was harvested, flushed with cold PBS to remove luminal content, and opened longitudinally. The tissue was cut into 3-5 mm pieces, washed thoroughly in cold PBS and incubated on ice in 5 mM EDTA for 10 min. After EDTA removal, intestinal pieces were vigorously vortexed for 3 min in cold PBS to release crypts. This process was repeated twice, and the third and fourth fractions (enriched in crypts) were pooled, filtered through a 70-µm cell strainer to remove remaining villi, and centrifuged at 1,000 RPM for 5 min. Pelleted crypts were washed in Advanced DMEM/F12 (#12634010, Thermo Fisher Scientific) and centrifuged again. The final pellet was resuspended in ice-cold BME (Cultrex #3532-010-02, BIO-TECHNE) and plated as domes, then incubated at 37°C. Organoids were cultured in IntestiCult™ Organoid Growth Medium (#06005, STEMCELL Technologies), hereafter referred as ENR medium. Organoids were passaged every 4-7 days by mechanical dissociation, and medium was refreshed every 2 days.

For bead insertion, organoids were mechanically dissociated during culture passaging, centrifuged, and then mixed with PAAm beads. The mixture was resuspended in ice-cold BME (Cultrex #3532-010-02 BIO-TECHNE), plated as domes, and incubated at 37°C.

### Immunofluorescence staining

*Cell immunostaining* - Cells were fixed with 4% paraformaldehyde (#15714, Electron Microscopy Science) for 10 min at room temperature, washed in PBS, permeabilized with 0.5% Triton 100-X for 10 min at room temperature, and blocked with 10% Foetal Calf Serum in PBS (blocking buffer) for 1 hour at room temperature. Primary antibodies were diluted in blocking buffer and incubated with the cell samples overnight at 4°C. Alexa Fluor-conjugated secondary antibodies were diluted at 1/200 in blocking buffer and incubated for 1 hour at room temperature. Phalloidin was added together with secondary antibodies. After each incubation step, samples were washed four times with PBS. Finally, samples were mounted in DABCO or home-made Mowiol antifade mounting medium.

*Organoid immunostaining* - At day 7, organoids were fixed using 4% paraformaldehyde for 30 min, then washed 3 times in PBS for 10 min each. Fixed organoids were stained with phalloidine-Atto647 (#65906) overnight, washed 3 times in PBS for 10 min each, and incubated in Hoechst 33342 for 15-30 min to stain nuclei. Immunostained organoids were mounted in home-made Mowiol solution.

Antibodies that have been used in this study are rabbit anti-atypical Protein Kinase C (#sc-216, Santa Cruz), rabbit anti-phospho-Myosin Light Chain-2 (Ser-19) (#3671, Cell Signalling Technologies), mouse anti-Paxillin (#B312405, BD transduction), rabbit anti-Paxillin phosphoTyr118 (#44-722G, Thermofisher Scientific), mouse anti-zyxin (#ab58210, Abcam), mouse anti-Vinculin (#14-9777-82, eBioscience), rabbit anti-β-catenin (#C2206, Sigma), donkey anti-rabbit Alexa488 (#A21206, Invitrogen), donkey anti-rabbit Alexa568 (#A10042 Invitrogen), donkey anti-rabbit Alexa647 (#A31573, Invitrogen), donkey anti-mouse Alexa488 (#A21202, Invitrogen), donkey anti-mouse Alexa568 (#A10037, Invitrogen), donkey anti-mouse Alexa647 (#715-605-151, Jackson ImmunoResearch). DAPI (4’,6-Diamidino-2-Phenylindole, Dihydrochloride, #D1306, Thermofischer scientific), Phalloidin-TRITC (#P1951, Sigma), Phalloidin-Atto-647N (#65906, Sigma).

### Image acquisition

Confocal images for bead size measurements were acquired with an inverted Zeiss LSM510 Meta confocal microscope equipped with C-Apochromat 40x/1,2 W objective. High-resolution confocal images for bead deformation analyses and immunostained cells or organoids were acquired using an inverted Zeiss LSM880 confocal microscope equipped with the C-Apochromat 40x/1,2 W autocorr M27 objective. Z-stacks spanning the epithelial architecture (i.e. cyst or organoid) were acquired for all samples. The z-step size was set to match the optimal sampling for the bead fluorescence signal (fluorescein-O-methacrylate) corresponding to 0.4333443 μm.

### Image processing for bead size measurements

Beads size was determined from maximum-intensity projections of the z-stack images. The resulting images were binarized, and watershed segmentation (ImageJ) was applied to separate contacting beads. The Feret’s diameter (maximum caliper diameter) of each bead was then measured using the Analyze Particles function in FiJi.

### Image processing for 3D bead deformation measurements

Bead deformation was analyzed using a custom Python pipeline. To reconstruct the 3D bead shape, image segmentation was performed on confocal z-stacks using the fluorescein-O-methacrylate signal, which uniformly labelled the entire bead volume. We found that the shape of beads that were not in contact with any cells appeared systematically flattened by a factor of approximately 1.038 in z direction when using DABCO mounting medium. We thus corrected all bead z stacks by this factor before quantifying the bead shape. Briefly, z-stacks were smoothed using a Gaussian blur (σ = 0.25μm). A defined number of rays were generated from the bead centre toward the exterior, uniformly distributed in latitude and longitude. Along each ray, fluorescence intensity was measured at regular distance intervals. Because the measurement point did not always coincide exactly with voxel positions, intensity values for each measurement points were obtained by linear interpolation. Since the beads were uniformly stained, fluorescence intensity sharply decreased at the bead surface, following a sigmoidal function. For each ray, the bead surface position was determined by fitting the intensity profile with an error function (the integral of a Gaussian function). The combination of all fitted surface positions was used to reconstruct the bead surface as a discretized grid. The initial bead center was estimated by hand. This was used to obtain a first estimate of the surface, which was then analyzed using spherical harmonics to refine the bead barycentre. In this way, in total four iterations were used to progressively refine the bead barycentre position. Once the bead surface was reconstructed, fluorescence intensity values from the different imaging channels were calculated at defined positions relative to the bead surface. In all plots, local bead deformation is quantified as the local radius deviation from the average bead radius, i.e. the integral of the radius deviation over the bead surface is zero.

In some cases, in particular for the beads embedded in organoids, we realized that with this method, we obtained substantial radius deviations also in parts of the bead that were not in contact with any cell. We hypothesized that this was because the barycentre of the deformed bead does not necessarily correspond to the barycentre of the undeformed bead. Thus, to find the material point of the deformed bead that corresponds to the center of the undeformed bead, we applied a midpoint correction to the beads embedded in organoids. To this end, we fitted a sphere to the cell-free surface of the deformed bead, varying its radius and midpoint. The cell-free surface was marked using a mask, and we verified that (i) varying the mask shape did not substantially affect results and (ii) the midpoint correction did not substantially affect the results for the embedded beads.

### Correlation analysis

To quantify the relationship between protein fluorescence distribution and bead deformation, 3D fluorescence datasets were analyzed using a surface-based projection approach. Briefly, grayscale image z-stacks corresponding to the protein-of-interest channel were extracted and projected onto the bead surface using a spherical coordinate framework, generating spatially resolved fluorescence intensity distributions aligned with the bead geometry. To enable comparison between samples, fluorescence intensities were normalized to the maximum intensity measured within each bead, resulting in relative intensity values ranging from 0 to 1.

For quantitative analysis, relative protein intensity and relative radial deformation (defined as deviations from an ideal spherical shape) were discretized using linear binning. One-dimensional histograms were generated for each parameter, and a two-dimensional joint histogram was computed to quantify their spatial co-distribution at the bead interface. To determine whether the observed correlations deviated from random association, an uncorrelated reference distribution was calculated as the outer product of the marginal distributions of fluorescence intensity and bead deformation. This provided a reference distribution assuming statistical independence between the two variables. The experimentally measured joint distribution was then compared with this uncorrelated reference distribution. The difference map (measured distribution minus uncorrelated distribution) was calculated and displayed using a diverging colormap, allowing the identification of enriched or depleted combinations of protein fluorescence intensity and bead deformation states.

### Force inference and FEM simulations

We aimed to directly infer the forces exerted by cells on the bead surfaces by inverting a linear elastic problem [34,57]. However, with our current experimental protocol, the unstressed reference shape of the beads is not available. Together with the fact that the inversion of the linear elastic problem is underdetermined, we found that results of the inversion depended on regularization parameters, which means that they could not be trusted. Future work will allow us to obtain snapshots not only of the deformed bead, but also the undeformed bead. Together with the information of the nanoparticles, this will turn the elastic problem into a properly conditioned one. Thus, so far, for illustrative purposes, we demonstrated the effect of certain types of applied force patterns on the deformed bead shape.

To solve the bead deformation resulting from some applied force pattern, we solved the following force balance problem:

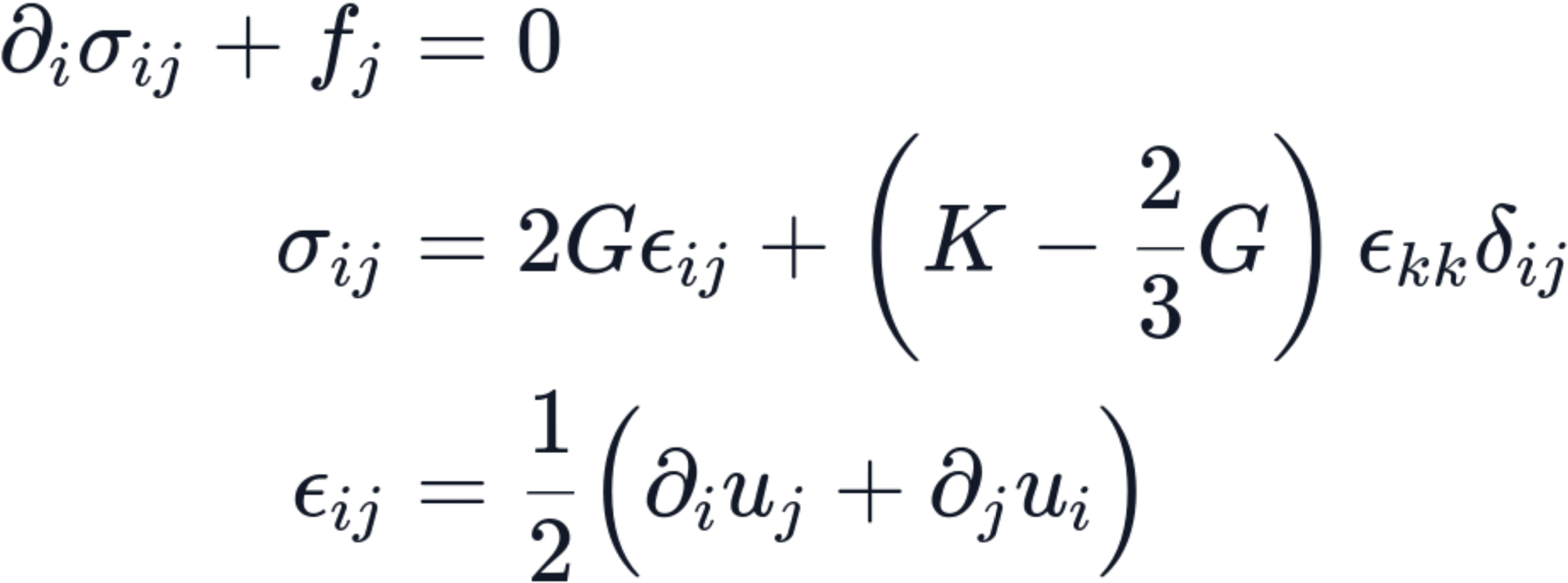

Here, *σ_i_*_j_ is the stress tensor, *f*_j_ is the bulk force distribution, *ε_i_*_j_ is the strain tensor, *G* is the shear modulus, *K* is the bulk modulus, and *u_i_* is the displacement field. These equations describe how stresses are transmitted across the bead through elastic coupling. Thus, our FEM solutions will include any potential non-local effects that may be created by such stress transmission.

We used Neumann boundary conditions, i.e. we imposed the boundary tension *T*_j_:

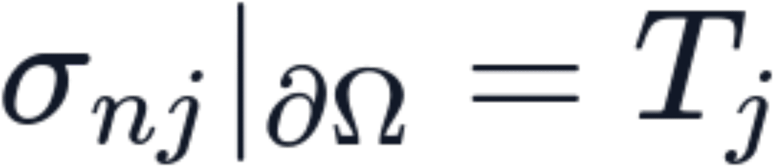

where *∂Ω* denotes the bead surface. For a spherical bead with radius 1, we set *T*_j_ to a localized normal force, 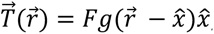, or a localized tangential force, 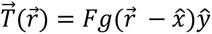. Here, *F* is a factor scaling the force and 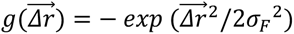 is a Gaussian with standard deviation *σ_F_*. We generated random normal forces by setting 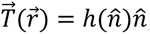, where 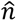 is the locally outward-pointing normal on the sphere surface and 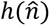 is created from a random superposition of spherical harmonics with power spectrum 1/(1 + *l*[*l* + 1]/*l*_0_^2^), setting the modes with *l* = 0 (homogeneous normal force) and *l* = 1 (creating net unbalanced forces on the sphere) to zero.

Some of these boundary conditions could end up applying a finite net force 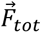 and/or net torque 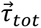 on the bead, acting contrary to overall force and torque balance. We thus defined the bulk force density by homogeneously spread-out, compensating force and torque:

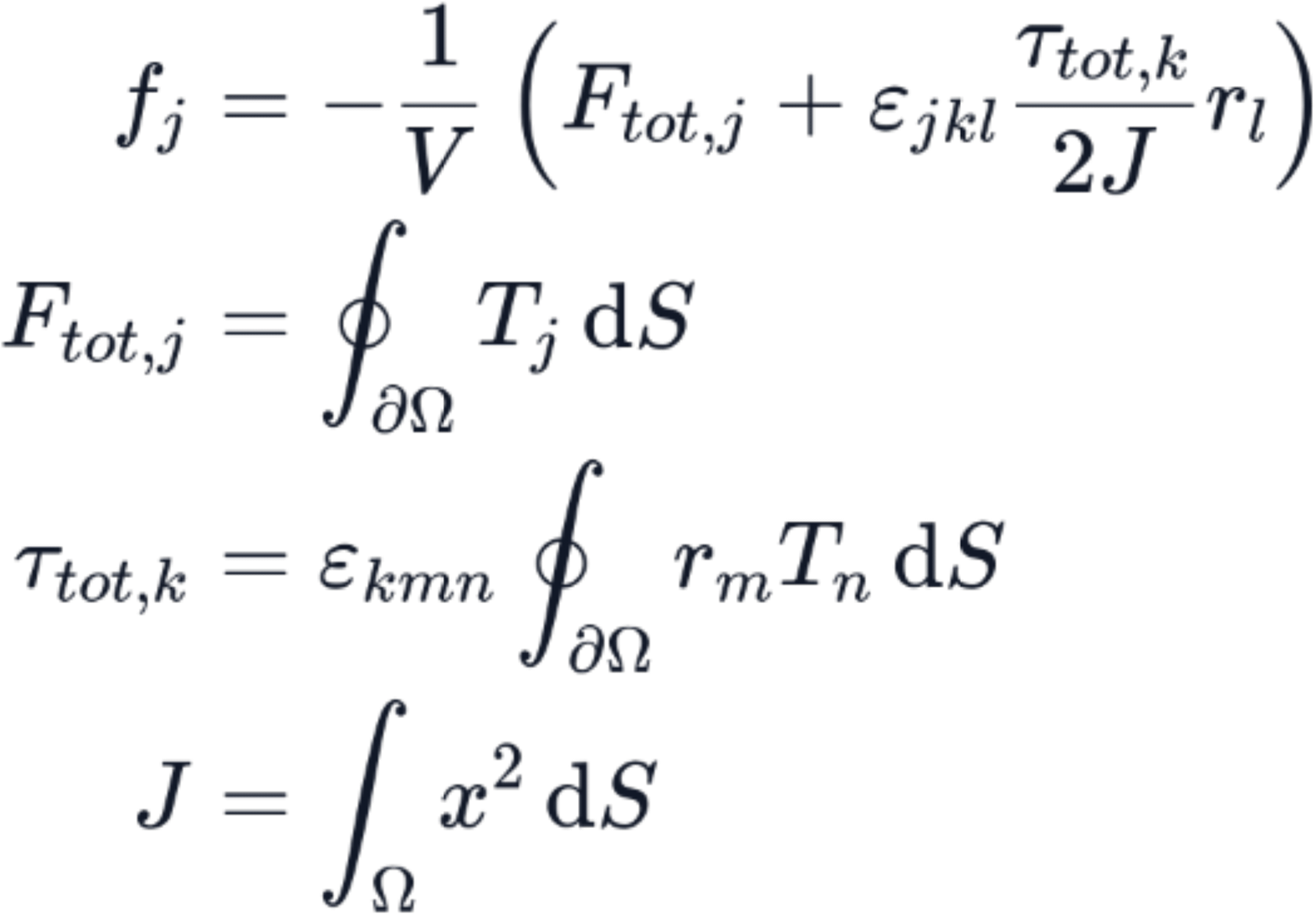

Here, *ε*_j*kl*_ is the Levi − Civita Symbol and *J* is the moment of inertia of the sphere.

We implemented the solution of this linear elastic problem in FEniCSx, version 0.11. Translational and rotational invariance of the problem was addressed by providing the corresponding six-dimensional nullspace to the solver.

We used as parameters *G* = 1 and *K* = 100 in dimensionless units. Note that radius deformation was measured with respect to the center of mass of the deformed bead, because this was the way this quantity was defined in the experiments. This explained the apparent positive displacement of the bead surface in places opposite to the force application.

### Statistical analysis

Statistical analysis was performed with R. The comparison of the bead diameters between soft and stiff beads was performed using the Mann-Whitney U test. The comparison between partially and completely inserted beads was performed with a Student’s t-test. The comparison between stiffnesses of soft and stiff beads was assessed using the Mann-Whitney U test. The comparison between bead deformations was performed using the Mann-Whitney U test. All statistical tests were performed after verifying the assumptions of normality (Shapiro-Wilk test) and homogeneity of variances (Levene’s test), when applicable. The comparison between the proportions of bead insertions depending on the coating condition was performed using a proportion test (test of equal proportions) with Yate’s correction. Three levels of statistical significance are distinguished: * p = 0.05; ** p = 0.01; *** p = 0.001, **** p < 0.001.

Unless otherwise specified in the Figure legend, all the experiments were performed with N = 3 independent replicates, with multiple technical replicates per experiment. The value of n corresponds to the total number of analyzed replicates pooled across the 3 N independent experiments.

### Code availability

Home-made codes and pipelines for bead segmentation and FEM simulations are available on GitHub (https://github.com/mmerkel/microbeads; https://github.com/mmerkel/microbeads_fem).

## Supplementary Materials

Supplementary Figures 1-4

