## Supplemental Files for "A non-invasive approach to study localized forces in 3D epithelia"

### Supplementary Materials

### Supplementary Figures 1-4

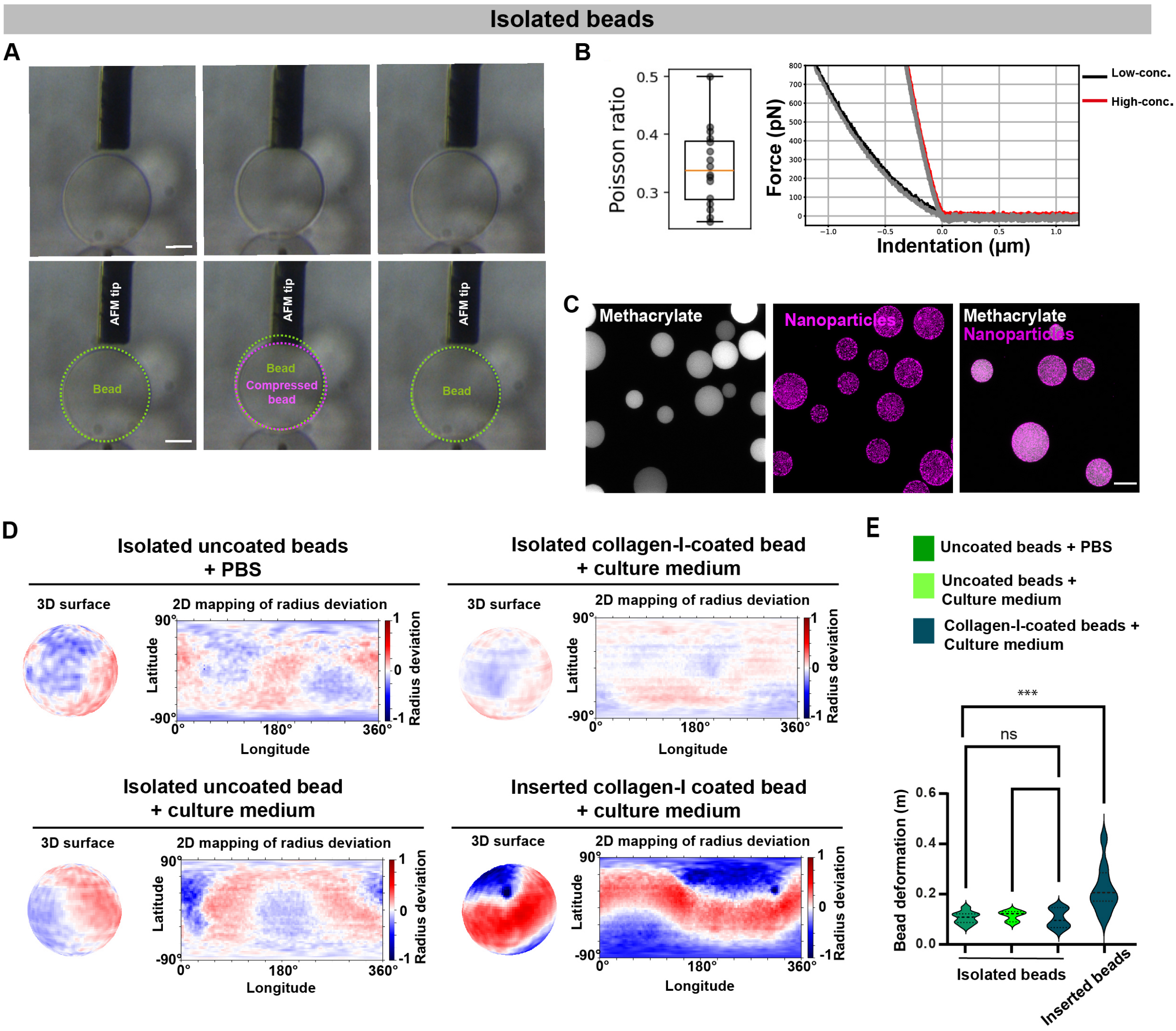


Gouirand et al., Supplementary Figure 1

**Supplementary Figure 1: Isolated PAAm beads are nearly incompressible and stable in different conditions**

**A,** Side view of a PAAm bead compressed by a platform cantilever. The green dotted circles outline the bead's initial contour (left), while the black dotted circle shows its deformed profile under a 200 nN applied force (middle). The vertical compression induces a horizontal enlargement, demonstrating the bead's incompressibility. Upon cantilever retraction and force release, the bead reverts to its initial shape (right), as expected for a fluid material. Scale bar, 20 µm. **B**, *Left,* box plot of the Poisson ratio of the PAAm beads assessed by measuring optically the lateral deformation while compressing the PAAm bead with the AFM cantilever (n=15 measurements over 10 beads). *Right,* representative AFM force-indentation curves showing that the low-concentration (black) and high-concentration (red) beads are purely elastic. The approach (black, red) and retract curves (grey) are coincident, indicating there is no viscous dissipation. **C**, Confocal analysis showing the distribution of fluorescein-O-methacrylatelabelled PAAm beads and far-red- carboxylate-modified nanoparticles PAAm beads. Scale bar, 20µm. **D**, 3D reconstruction of the bead surface (left) and 2D projection representation (right), of beads under following conditions: isolated uncoated beads in PBS (top panels left), isolated uncoated beads in culture medium (bottom panels left), isolated collagen-I-coated beads in culture medium (top panels right), and collagen-I-coated beads inserted within Caco-2 cysts (bottom panels right). Positive radius deviations are shown in red, and negative radius deviations in blue. **E**, Quantification of PAAm bead deformation under the following conditions: isolated uncoated beads in PBS, isolated uncoated beads in culture medium, isolated collagen-I-coated beads in culture medium, and collagen-I-coated beads inserted within Caco-2 cysts. Violin plots show statistical analysis of the bead deformation (in µm) under each condition. n = 8 isolated uncoated beads in PBS, n=3 isolated uncoated beads in culture medium, n = 10 isolated collagen-I-coated beads in culture medium, and n = 10 collagen-I-coated beads inserted within Caco-2 cysts. Mann-Whitney test, ***p< 0,01.

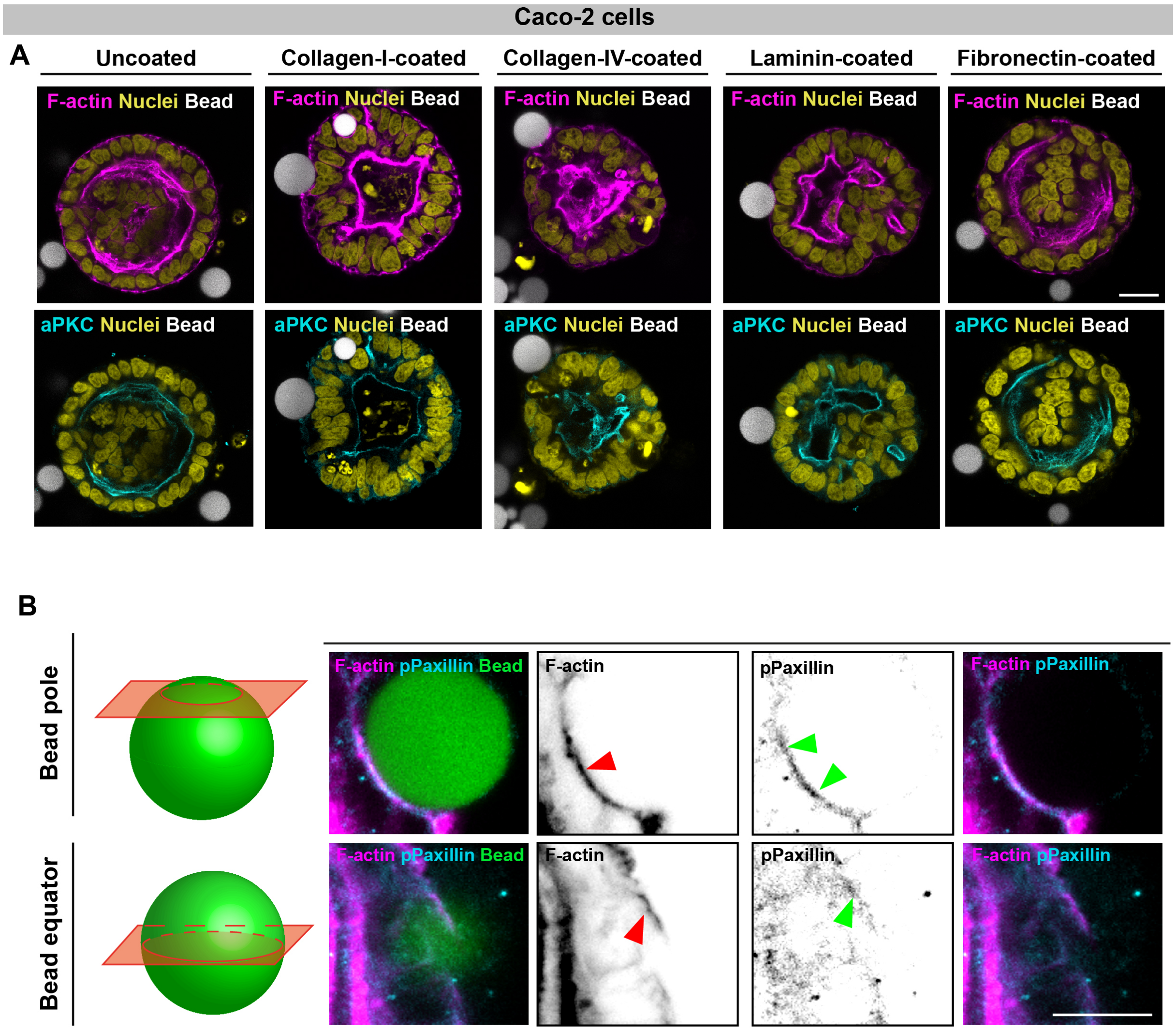


Gouirand et al., Supplementary Figure 2

**Supplementary Figure 2: Insertion of ECM-coated PAAm beads does not alter Caco-2 cyst organization and leads to phospho-paxillin enrichment at the bead-cell interface**

**A,** Confocal analysis of Caco-2 cysts containing uncoated beads (control), collagen-I-coated beads, collagen-IV-coated beads, laminin-coated beads, or fibronectin-coated beads. In the top panel, cysts were stained for F-actin with phalloidin (magenta), nuclei with DAPI (yellow), and beads are shown in grey. Bottom panels show the corresponding regions stained for aPKC (cyan), with nuclei (DAPI, yellow) and beads (gray). Scale bar: 20µm. **B,** Left, schematic defining the pole and equatorial imaging planes used for bead analysis. Right, confocal images of a collagen-I–coated bead interacting with cells at the pole and equatorial planes (F-actin: magenta; pPaxillin cyan; bead: green). Grayscale images display individual fluorescence channels, and merged images illustrate their spatial organization. Arrowheads indicate regions of F-actin (red) and pPaxillin (green) enrichment or depletion at the bead-cell interface. Scale bar: 10 µm.

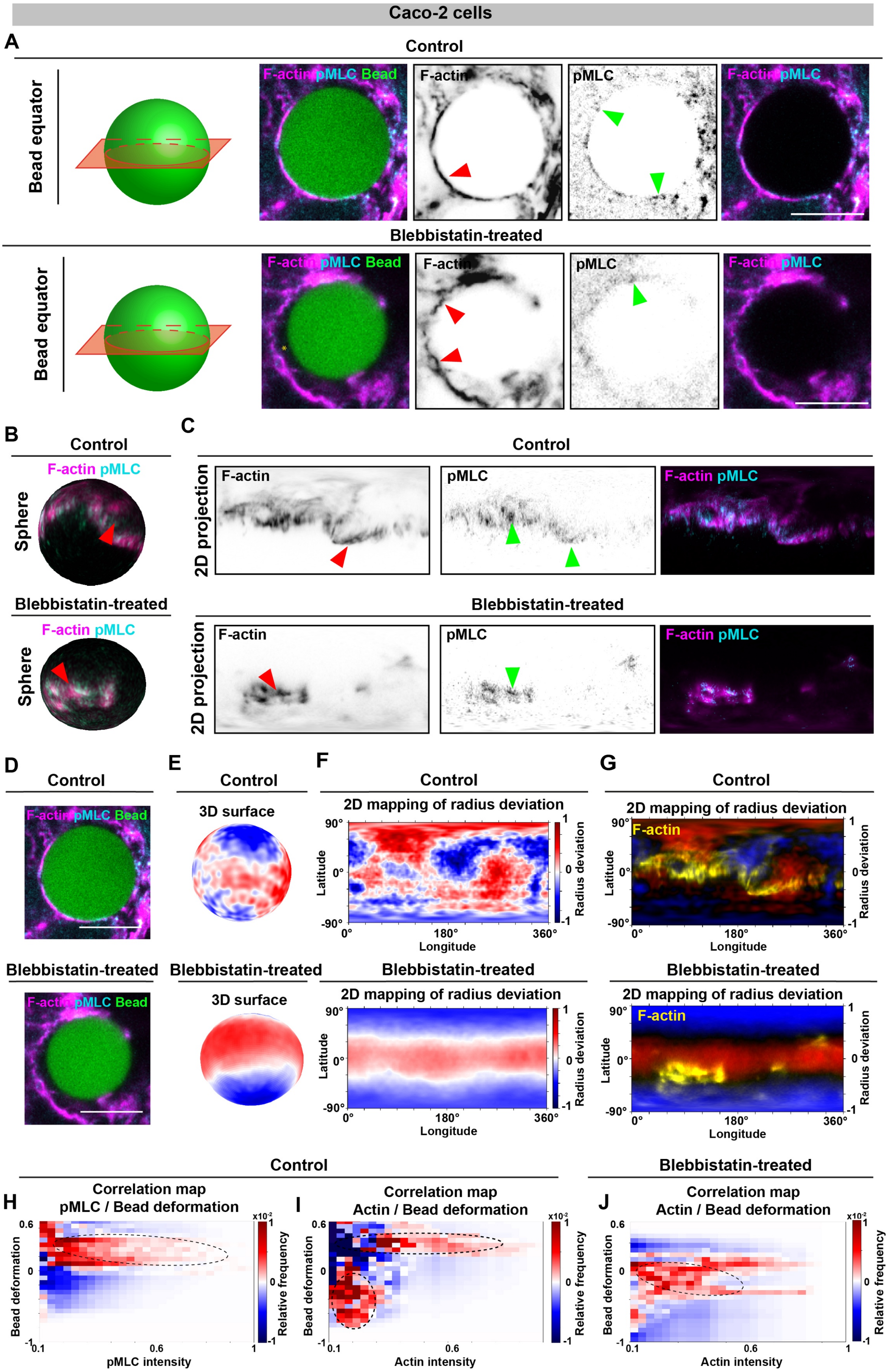


Gouirand et al., Supplementary Figure 3

**Supplementary Figure 3: Inhibition of contractility decouples bead deformation from actomyosin organization at the bead-cell interface**

**A**, Left, schematic defining the equatorial imaging plane used for bead analysis. Right, confocal images at the equatorial planes of collagen-I–coated beads interacting with control cells (top panel) or blebbistatin-treated cells (bottom panel) (F-actin: magenta; pMLC: cyan; bead: green). Grayscale images show the individual fluorescence channels, and merged images illustrate their spatial organization. Arrowheads indicate regions of F-actin (red) and pMLC (green) enrichment or depletion at the bead-cell interface. Scale bar: 10 µm. **B-C**, 3D spherical mapping (**B**) and 2D projection representation (**C**) of pMLC (cyan) and F-actin (magenta), and their merged signals at the bead-cell interface in control cells (top panel) and blebbistatin-treated cells (bottom panel). Arrowheads indicate regions of F-actin (red) and pMLC (green) enrichment at the bead-cell interface. **D**, Confocal images at the equatorial plane of collagen-I–coated beads interacting with control cells (top panel) or blebbistatin-treated cells (bottom panel) (F-actin: magenta; pMLC: cyan; bead: green). **E-F,** 3D reconstruction of the deformed bead surface (**E**) and 2D projection representation (**F**) for control cells (top panel) and blebbistatin-treated (bottom panel) cells. Positive radius deviations are shown in red, and negative radius deviations in blue. **G**, Overlay of the 2D projection representation and actin signal (yellow) of control cells (top panel) and blebbistatin-treated cells (bottom panel). **H-J,** Representative correlation maps of the bead deformation (left, y-axis), pMLC intensity in control condition (**H**, x-axis), actin intensity in control condition (**I**, x-axis), actin intensity in blebbistatin-treated condition (**J**, x-axis) and frequency of occurrence (right, y-axis). High frequency is shown in red, and low frequency in blue.

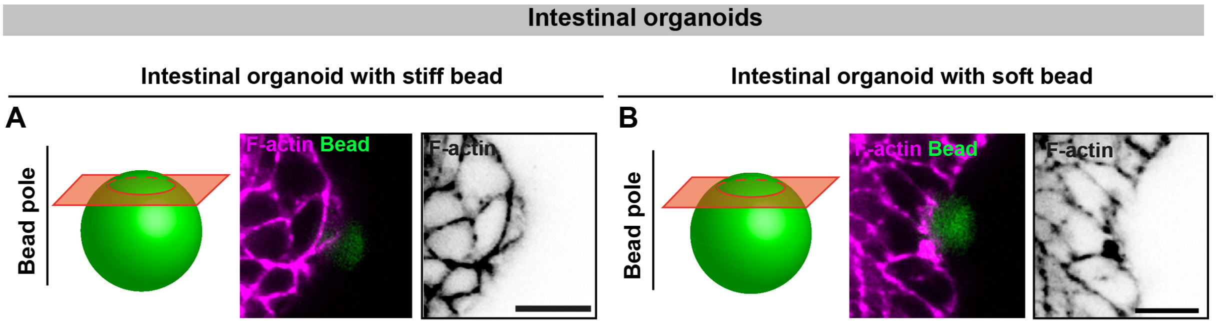


Gouirand et al., Supplementary Figure 4

**Supplementary Figure 4: Bead pole view of its insertion into the crypt-like domain of intestinal organoids**

**A**, Left, schematic defining the pole imaging plane of beads. Right, confocal analysis at the pole plane of a collagen-I–coated stiff bead interacting with cells (F-actin: magenta; bead: green). Scale bar: 10 µm. **B**, Left, schematic defining the pole imaging plane of beads. Right, confocal analysis at the pole plane of a collagen-I–coated soft bead interacting with cells (F-actin: magenta; bead: green). Scale bar: 10 µm.
